# A human-specific, concerted repression of microcephaly genes contributes to radiation-induced growth defects in forebrain organoids

**DOI:** 10.1101/2024.06.27.600564

**Authors:** Jessica Honorato Ribeiro, Emre Etlioglu, Jasmine Buset, Ann Janssen, Hanne Puype, Lisa Berden, André Claude Mbouombouo Mfossa, Winnok H. De Vos, Vanessa Vermeirssen, Sarah Baatout, Nicholas Rajan, Roel Quintens

## Abstract

Prenatal radiation-induced DNA damage poses a significant threat to normal brain development, resulting in microcephaly which primarily affects the cerebral cortex. It is unclear which molecular mechanisms are at the basis of this defect in humans as the few mechanistic studies performed so far were done in animals. Here, we leveraged human embryonic stem cell-derived forebrain organoids as a model for human corticogenesis. Organoids were X-irradiated with a moderate and a high dose at different time points, representing very early and mid corticogenesis. Irradiation caused a dose- and developmental-timing-dependent reduction in organoid size, which was more prominent in developmentally younger organoids. This coincided with a dose-dependent canonical p53-DREAM-dependent DNA damage response (DDR), consisting of cell cycle arrest, DNA repair and apoptosis. The DDR was delayed and less pronounced in the older organoids. Besides the DDR, we observed radiation-induced premature differentiation of neural progenitors and changes in metabolism. Importantly, our transcriptomic analysis furthermore demonstrated a concerted p53-E2F4-dependent repression of primary microcephaly genes. We found that this was a human-specific feature, as it was not observed in mouse embryonic brains or primary mouse neural progenitor cells. Thus, human forebrain organoids are an excellent model to investigate prenatal DNA damage-induced microcephaly and to uncover potentially targetable human-specific pathways.

## 1. Introduction

The development of the human brain depends on a delicate balance among neural stem and progenitor cell maintenance, proliferation, differentiation, and migration, in a spatiotemporal-specific manner (1). When these processes are disrupted, normal development can be drastically compromised, potentially resulting in neurodevelopmental disorders (2,3). A common feature of these disorders is microcephaly, a defect that primarily affects the cerebral cortex and is associated with intellectual disability, especially when it occurs at birth (primary microcephaly) (4,5). Microcephaly primary hereditary (MCPH), a non-syndromic genetic form of primary microcephaly, is the most well-characterized subtype, for which 30 genes have been identified as causative and further referred to as “MCPH genes” (6,7). These genes are involved in centrosome biogenesis and DNA repair pathways, with patients often displaying genomic instability due to DNA damage accumulation and radiosensitivity (8,9). This underscores the particular sensitivity of the developing brain to DNA damage, which is further illustrated by the fact that DNA repair deficiency syndromes often display microcephaly as one of the most prominent features (8,10). Also, environmental exposures that can lead to microcephaly are prenatal irradiation or Zika virus infections, both of which result in DNA damage accumulation in neural progenitors (NPCs) (11,12).

In the case of ionizing radiation (IR) exposure, the risk is particularly significant when it occurs during early corticogenesis between gestational weeks 8 to 15, and to a lesser extent between gestational weeks 15 to 26 (11,13). The latter reinforces the differences in sensitivity observed across developmental stages. One possible explanation for this vulnerability is the lower apoptosis threshold displayed by NPCs in response to DNA damage when compared with progenitors generated at later developmental stages (14). Apoptosis is one of the possible outcomes of the DNA damage response (DDR), a tightly controlled set of mechanisms to counteract genotoxic stress. The DDR typically involves cell cycle arrest, allowing cells to repair the DNA via different pathways that depend on the type and the number of lesions (15). In case of insufficient repair, cells may undergo apoptosis, senescence or differentiation as ways to escape malignancy (16,17). One of the key regulators of the DDR is p53, a transcriptional activator of genes involved in cell cycle arrest, DNA repair and apoptosis. Besides direct transactivation of these genes, p53 also indirectly represses cell cycle genes via the activation of CDKN1A (p21/CIP1/WAF1) (18,19). p21 leads to hypo-phosphorylation of RB-related pocket proteins p107 and p130. In such state, p107/p130 can bind to other proteins and form the DREAM complex (MuvB core complex, E2F4-5/DP, and p130 or p107), which regulates transcriptional repression through binding of the repressor E2F4 or -5 transcription factors (TFs) on E2F binding sites within gene promoters (18,20).

The current known mechanisms driving DNA damage-induced microcephaly have primarily been studied in mouse models, which have offered valuable insights into how normal brain development can be disrupted (21–23). Excessive DNA damage during early neurogenesis has been shown to severely affect NPCs populating the ventricular zone and subventricular zone of the developing cortex, culminating in NPC pool depletion (24–27). Among the underlying mechanisms, are the shift from proliferative to differentiative cellular divisions resulting in premature differentiation (28). Also G1 lengthening and prolonged mitosis leading to precocious neurogenesis or apoptosis has been observed (3,4,29). Additionally, mitotic defects and centrosomal aberrations that disrupt proper cell division have also been seen (21). While these processes are extremely relevant, the primary mechanism implicated in DNA damage-induced microcephaly, including that resulting from IR exposure, involves the activation of p53-dependent responses (23,30–33). In the mouse cortex, hyperactivation of p53 has been shown to induce widespread apoptosis of NPCs, disrupting normal brain development and contributing to microcephaly (23,24,27,33). Despite the valuable information gathered from these mouse models, they often did not fully recapitulate the human phenotype. Examples are mouse models of critical human MCPH genes such as *ASPM*, *WDR62* or *CDK5RAP2*, which showed only mild effects on neurodevelopment compared with patients with orthologous mutations (34–38). Although most of the aforementioned mechanisms have also been reported in mouse models of radiation-induced microcephaly (39), little is known about the mechanisms underlying irradiation of human models. A few previous studies have investigated short-term effects of radiation on human brain organoids. However, these studies focused on the early DDR and were not conducted in the context of microcephaly (40,41).

Therefore, there is a need for a better understanding of the disease etiology following IR exposure in humans. The available data on the safety of radiation exposure during pregnancy are based on a limited number of animal studies and studies on atomic bomb survivors (42). Based on the latter, the radiation protection systems follow the “as low as reasonably achievable” principle. However, because of the large uncertainties about the assumed risks (including also cancer) to the unborn child, in current clinical practice this comes with the consequence that even for extra- abdominopelvic cancers, treatment of pregnant women with radiotherapy is mostly contraindicated and postponed until after delivery (42). Consequently, less than 2% of pregnant cancer patients receive radiotherapy while in the general patient population this number amounts to around 50% (43). This may be problematic when RT is essential for primary treatment, resulting in suboptimal treatment of the patients which can be harmful for both patient and child (44).

In this study, human embryonic stem cell (hESC)-derived forebrain organoids (45) were used to model the early and mid-stages of cortical development and investigate effects of moderate (0.5 Gy) and high doses (2 Gy) of IR (X-rays). Our findings showed a dose- and developmental-timing-dependent reduction in organoid size, resembling microcephaly. This could be explained by increased apoptosis and premature differentiation of NPCs in irradiated organoids. Furthermore, our study uncovered a coordinated p53-E2F4/DREAM-dependent downregulation of MCPH genes that could explain part of the growth defects observed after irradiation. Importantly, this is a human-specific feature of the developing brain’s radiation response as it was not seen in mouse embryonic brains or NPCs.

## 2. Results

### hESC-derived forebrain organoids resemble early human cortical development

First, organoid size was monitored during culture which indicated a constant growth (Figure 1A). In order to assess our organoid cultures as a proper model for human brain development, we next characterized them at four developmental time points: D14, D28, D56, and D70. These resemble early to mid-fetal brain development which is the most critical phase for radiation-induced neurodevelopmental defects (39). The presence of specific cell types was investigated using immunostaining for cellular markers (Figure 1B). The expression of the neural stem cell markers SOX2 and NES was observed throughout organoid development, while the early neuroectoderm progenitor and dorsal pallium marker PAX6 was observed from D28 onwards. The numbers of proliferating cells decreased, which coincided with an increase in cells expressing the immature neuronal marker DCX, present from D28 onwards, and mature neurons expressing TUJ1 which were also visible from D28 onwards, but showing a substantial increase by D56. Oligodendrocyte progenitor cells positive for OLIG2 appeared from D28 onwards with a sharp increase in their prevalence at D56, albeit that their total numbers were still limited.

**Figure 1.**
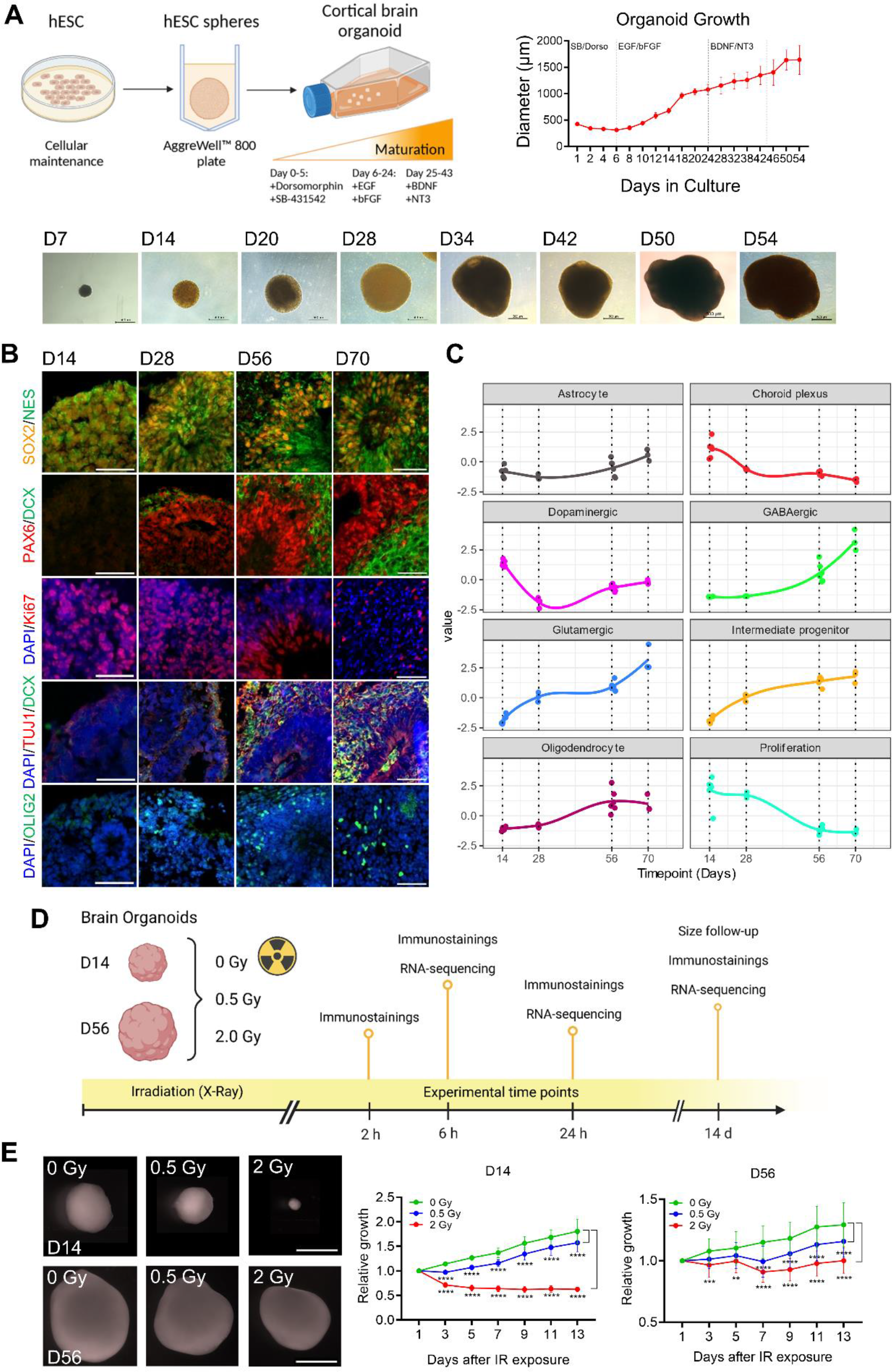
Forebrain organoid growth is dose- and time-dependently reduced by ionizing radiation (IR) exposure. Early human cortical development was modeled using forebrain organoids derived from embryonic stem cells (hESCs), following the methodology by Sloan et al., 2019. (A) Every organoid batch was generated using 600 microwells of an AggreWell™ 800 plate. Double SMADi inhibition with 5 µM Dorsomorphin and 10 µM SB-431542 was applied from Day 0 to Day 5, followed by exposure to 20 ng/ml EGF and bFGF from Day 6 to Day 24. Subsequently, 20 ng/ml BDNF and NT-3 were applied from Day 25 to Day 43. From Day 43, cell culture medium was free from growth factors. Organoid growth was followed up from Day 1 to Day 54, and the diameter of at least 10 organoids per time point was quantified using the ImageJ software. Scale bar = 500 µm. (B) Forebrain organoid characterization via immunohistochemistry revealed the expression of classical markers present throughout normal human cortical development: SOX2, Nestin, and PAX6 staining for neural progenitor cells, DCX and TUJ1 staining for post-mitotic neurons, OLIG2 for oligodendrocytes and oligodendrocyte precursors. Scale bar = 50 µm. (C) Deconvolution analysis based on single cell RNA-seq data (from (51)) showed the abundance of different cell types during organoid development. (D) Forebrain organoids were exposed to ionizing radiation (Sham-irradiated x 0.5 Gy x 2 Gy) at developmental Days 14 and 56. Samples were processed for immunostaining (2 h, 6 h, 24 h, 14 d), RNA-sequencing (6 h, 24 h, 14 d), and size follow up (14 d). (E) Brain organoid size was monitored every 2 days. 24 organoids were analyzed per condition. Scale bar = 1000 µm. Statistical analysis was performed using Two-way ANOVA test followed by Tukey’s test for multiple comparisons. **p< 0.001; ***p< 0.0004; ****p< 0.0001. See also Figure S1-S3.

Transcriptomic profiling of organoids at the same time points using bulk RNA-sequencing (RNA-seq), confirmed the developmental trajectory during organoid culture. The most pronounced changes occurred between D14 and D28 and between D28 and D56, while changes between D56 and D70 were overall more subtle (Figure S1A; Table S1). Gene ontology (GO) enrichment analysis indicated that genes enriched at later time points of organoid development were mostly involved in typical neurodevelopmental processes such as axonogenesis, synaptogenesis and synaptic transmission, reflecting neuronal differentiation and maturation (Figure S1B). We also observed increased expression of genes related to the mitochondrial electron transport chain, which was most evident between D28 and D56. This corresponds to the switch from glycolysis to oxidative phosphorylation that occurs during neurogenesis and neuronal maturation (46). Developmentally enriched genes were predicted to be mainly regulated by TFs like REST, SMAD4 and TP53, and the PRC2 members EZH2 and SUZ12, as evidenced by Enrichr analysis (Figure S1C). REST (also called Neuron-Restrictive Silencing Factor, NRSF) is a critical repressor of neuronal genes in non-neuronal tissues and neural stem and progenitor cells (47), which recruits the PRC2 complex to neuronal gene promoters via interaction with EZH2 and SUZ12 (48). The function of REST, EZH2 and SUZ12 as transcriptional repressors, yet being predicted to regulate developmentally enriched genes was consistent with their own downregulation during organoid development (Figure S2A). Their expression profiles are in line with those from other studies in human brain tissue and human brain organoids (Figure S2B-D). SMAD regulates neural differentiation and SMAD inhibition is often used to induce neural conversion of ESCs (49). SMAD4 itself is also reduced during organoid maturation (Figure S2B-D). The role of p53 in brain development is still poorly understood, although a small fraction of *Trp53* knockout mice develop exencephaly (50). In general, however, typical p53 target genes are downregulated during brain development, although we found previously that a subset of p53 targets are developmentally upregulated in the embryonic mouse brain (31). This is in contrast with *TP53*/*Trp53* itself, which is downregulated during brain and organoid development (Figure S2B-D).

Genes that were downregulated with time were mostly involved in processes such as DNA replication, DNA repair, the DNA damage response, and cell cycle regulation, especially during the early stages of organoid development. Later, between D56 and D70, genes involved in RNA processing and ribosome biogenesis became additionally downregulated (Figure S1D). Downregulated genes were predicted to be primarily regulated via E2F family TFs, especially E2F4, and the NFYA-B family (Figure S1E).

We deconvolved the cellular composition of the organoids in function of their maturation based on single cell RNA-seq data from forebrain organoids of similar developmental age cultured using the same protocol (51). This analysis revealed the presence of various cell types, including intermediate progenitors, neurons (dopaminergic, GABAergic, and glutamatergic), astrocytes, oligodendrocytes, and choroid plexus cells (Figure 1C). It indicated that with time, the number of proliferating progenitors and choroid plexus (ChP)-like cells were overall reduced. In contrast, cell types associated with later stages of brain development such as intermediate progenitors, neurons (both inhibitory and excitatory) and glial cell types (astrocytes, oligodendrocytes) were enriched during organoid culture (Figure 1C). These cells emerge at different developmental time points, resembling the proper human brain development, and correlate with the gradual reduction in proliferation observed. For instance, there is an increasing presence of post-mitotic neurons over time (Figure 1C). Examples of expression patterns of different markers during the course of organoid growth are indicated in Figures S1F-G. In alignment with human cortical development, NPC markers such as HMGA2 are predominantly seen during early stages of organoid development. On the other hand, markers such as SOX2 and Nestin (NES) exhibit sustained expression over time, as seen at the protein level, although with a gradual decline, with a more region-restricted expression (neural rosettes) as neurodevelopment progresses. Markers of intermediate progenitors (EOMES/TBR2, PPP1R17), outer radial glia cells (TNC), and postmitotic neuronal markers, such as TUBB3/TUJ1, and NEUROD6 emerge at later stages, as well as specific markers of specific neuronal populations like BCL11B/CTIP2 (deep layer neurons), RELN (Cajal-Retzius neurons), GAD1 and GAD2 (interneurons), oligodendrocyte precursor markers like OLIG1 and OLIG2, and astrocyte markers like ATP1B2. Overall, the gene expression profiles and protein levels of cellular markers observed during the development of our organoid cultures indicated their suitability as models for human corticogenesis.

### Forebrain organoid growth is dose- and time-dependently reduced by ionizing radiation (IR) exposure

Forebrain organoids were irradiated at D14 and D56 with X-ray doses of 0.5 Gy and 2 Gy. Samples were processed for immunostainings, RNA sequencing, and their size was monitored for 14 days after irradiation (Figure 1D). For the lower dose, organoid growth stagnated for three and seven days in organoids irradiated at D14 or D56, respectively, after which their growth resumed at a normal pace, although not reaching the same size as that of sham-irradiated controls (Figure 1E). Organoids irradiated with a high dose at D14 significantly reduced in size and did not recover (Figure 1E). In fact, 14 days after exposure these organoids were so small that they could not be used for subsequent immunostaining experiments. In contrast, 2-Gy irradiated organoids at D56 showed a reduction in size until day 7 but resumed their growth thereafter (Figure 1E). In addition to the reduction in size, morphological changes in neural rosettes within forebrain organoids were also observed after a 14-day follow-up. Organoids irradiated at D14 with a 0.5-Gy dose displayed rosette enlargement both in terms of perimeter and thickness accompanied by the loss of normal cellular architecture that consists of NPCs radially organized around the lumen (Figure S3A). However, D56 organoids irradiated with 0.5- and 2-Gy doses displayed a reduction in rosette size and thickness over time (Figure S3B). We believe that these contrasts in rosette morphology between D14 and D56 are related to differences in response to IR at distinct developmental time points, which will be further addressed in the following sections of this manuscript. Overall, organoids showed a time-, dose-, and developmental time-dependent reduction in size along with changes in neural rosette morphology after exposure to IR.

### Irradiation of forebrain organoids leads to the accumulation of DNA damage followed by a p53-dependent DNA damage response and premature neuronal differentiation

IR causes DNA damage which normally induces a DDR (8). To investigate the activation of radiation-induced DNA damage in forebrain organoids, immunostainings were conducted for phosphorylated histone H2AX (ɣH2AX) and p53-binding protein 1 (53BP1), which both localize to DNA double-strand breaks (DSBs). DSB foci formation could be observed at 2 h post-irradiation for both D14 and D56 organoids, and this effect was dose-dependent (Figure 2A,B). At 6 h following IR exposure, DSB foci number was reduced compared to 2 h but still significant when compared to the 6 h control for both developmental time points (Figure 2A,B). At 24 h, baseline levels of DSB foci were reached in D14 organoids (Figure 2A), while increased numbers were still present at D56 for both IR doses (Figure 2B). This reflects the enhanced ability of DNA repair by NPCs at early developmental stages compared to more differentiated cell types present at later time points (54). Additionally, they could indicate the high sensitivity of NPCs to damage, which will be discussed further.

**Figure 2.**
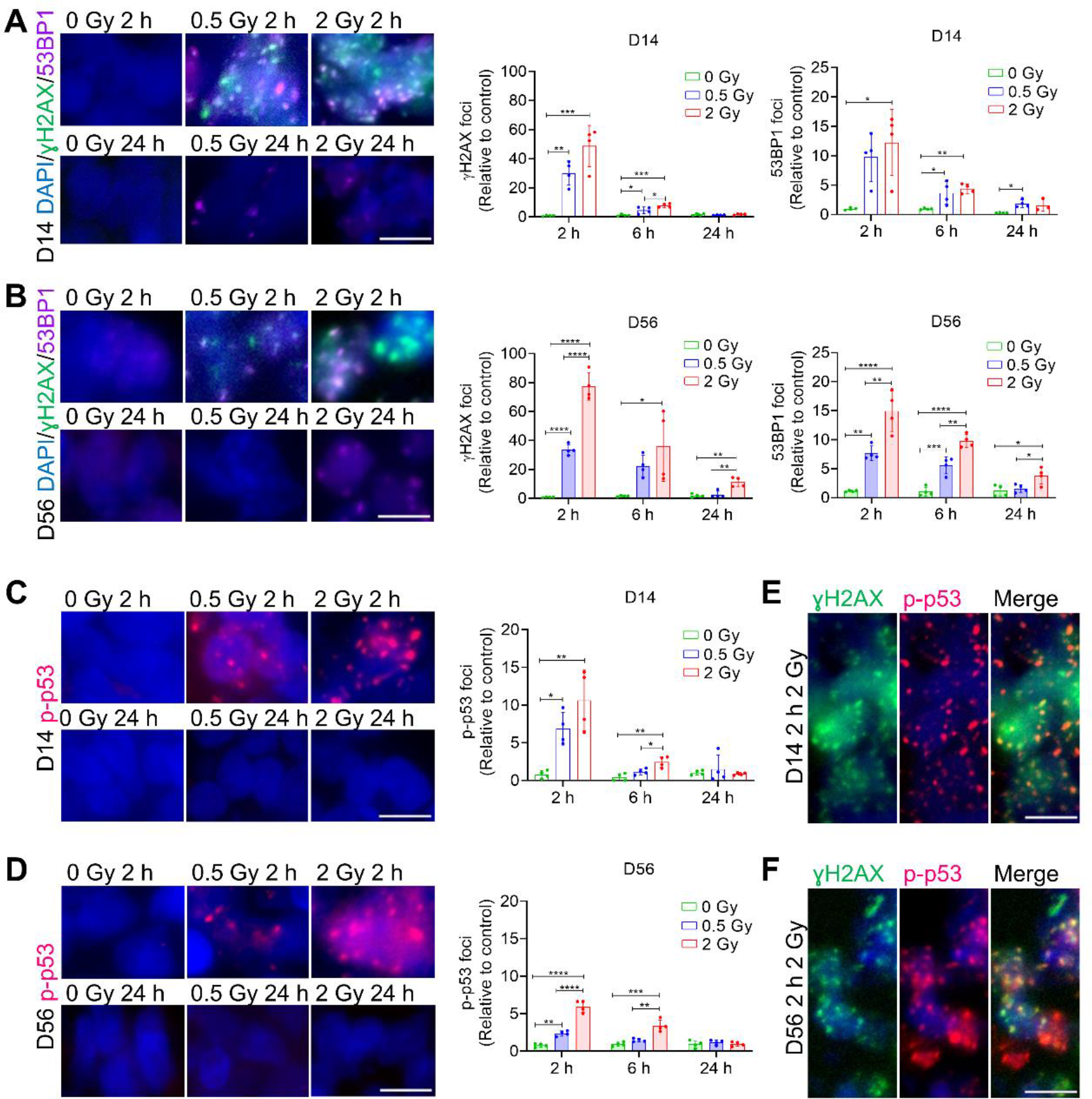
Ionizing radiation exposure induces double-strand breaks and p53 recruitment to DNA damage sites, in a dose- and developmental stage-dependent manner. (A, B) DNA damage induction was analyzed via immunostaining of DNA double-strand break (DSB) sensitive markers: phosphorylated histone H2AX (ɣH2AX) and p53-binding protein 1 (53BP1). (C, D) DNA damage response (DDR) was analyzed via immunostaining of phosphorylated p53 (p-p53). (A-D) Analysis were performed at 2 h, 6 h, and 24 h post-irradiation of D14 and D56 organoids. n = 4. Image analysis was performed using the ImageJ software. One-way ANOVA test followed by Tukey’s test for multiple comparisons was used, *p<0.04 **p< 0.005; ***p< 0.0009; ****p< 0.0001. (E, F) Co-localization of ɣH2AX and p-p53 foci 2 h post-exposure to a 2 Gy dose in D14 and D56 organoids. (A, B) Scale bar = 5 µm. (C, D) Scale bar = 10 µm. (E, F) Scale bar = 10 µm.

The DDR is regulated via phosphorylation of p53 (p-p53) which then acts as a TF to activate genes involved in processes such as cell cycle arrest, DNA repair and apoptosis (55). Staining for p-p53 showed that 2 h post-irradiation there was an increased number of p-p53 foci present at both developmental stages, which seemed more pronounced in D14 organoids (Figure 2C,D). By 6 h post-irradiation, p-p53 response had ceased, and by 24 h p53 was inactive (Figure 2C,D). Lastly, an overlap of ɣH2AX and p-p53 foci was observed in both irradiated D14 and D56 organoids (Figure 2E,F). The rapid p53 recruitment to DNA damage sites was described as a unique property of human p53, and has been shown to influence the choice of repair pathway (56).

Following DNA damage, p-p53 acts as a crucial regulator of the cellular response, primarily inducing cell cycle arrest (53). This temporary pause in cell division provides time for DNA repair mechanisms to correct damages to the DNA before the cell progresses through the cycle. Therefore, we proceeded to investigate cell cycle dynamics. For this, immunostaining for phosphorylated histone H3 (PH3), and phosphorylated Vimentin (pVim) was performed. PH3 and pVim stains for cells that are actively undergoing mitosis. The impact of IR exposure on cell division was obvious, 2 h post-irradiation both PH3- and pVim-expressing cells were significantly decreased (Figure 3A- D). This effect was even more pronounced in younger organoids, which displayed near zero dividing cells following IR exposure at both radiation doses (Figure 3A,B). At 6 h post-irradiation, mitotic cells were still absent in D14 organoids irradiated with 2 Gy, while in those irradiated with 0.5 Gy some dividing cells could be seen (Figure 3C). At 24 h, organoids irradiated with a 0.5 Gy dose seemed to have recovered their proliferative potential, while those irradiated with 2 Gy displayed only a few dividing cells at both developmental time points (Figure 3C,D). Fourteen days following IR exposure, cell cycle dynamics for D56 organoids appeared to be normal (Figure 3D). Overall, these findings suggest that exposure to DNA damage induced by IR promotes cell cycle arrest in human forebrain organoids.

**Figure 3.**
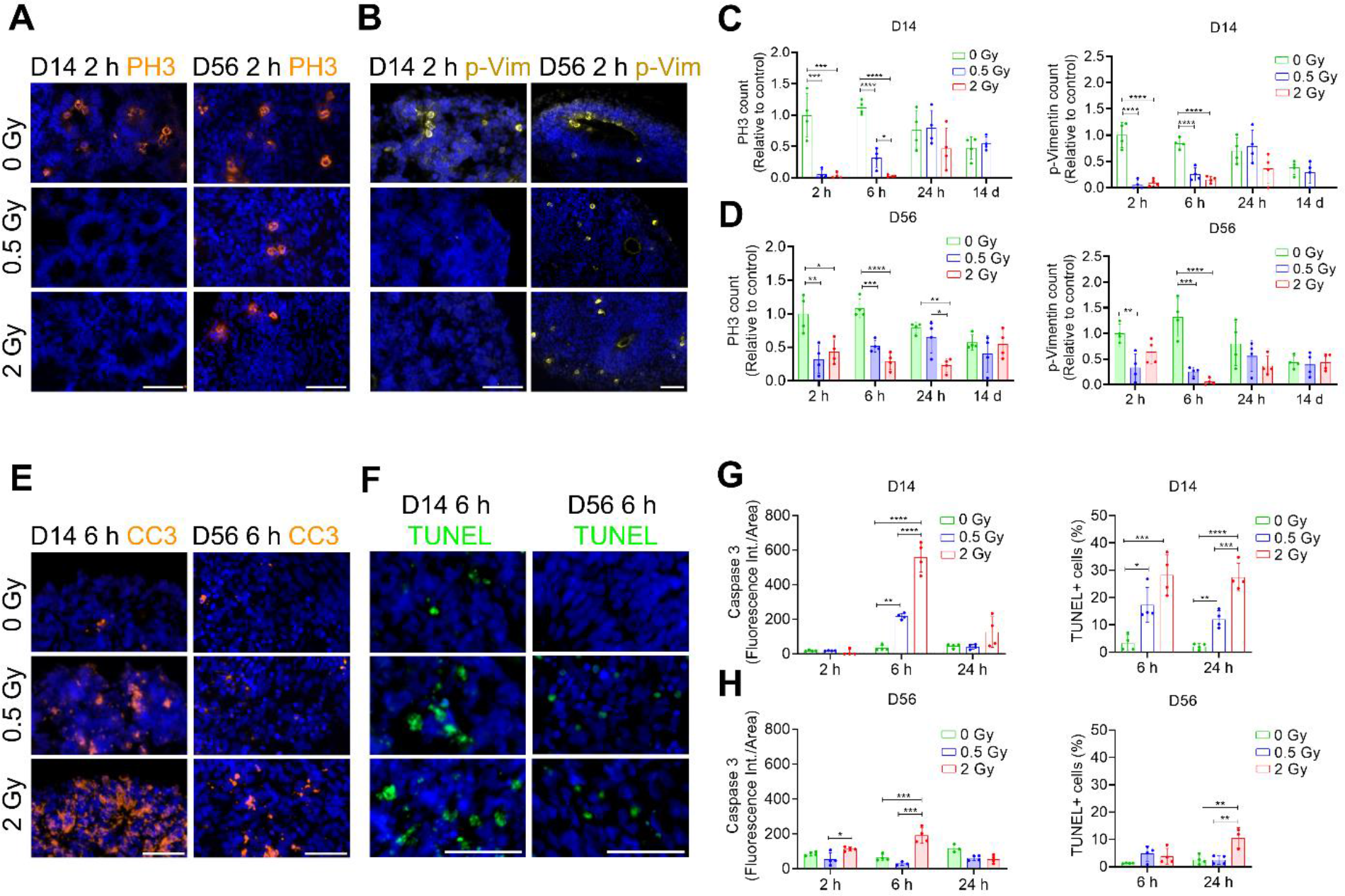
Ionizing radiation-induced DNA damage leads to cell cycle arrest and apoptosis in human forebrain organoids. Forebrain organoids were exposed to ionizing radiation (Sham- irradiated x 0.5 Gy x 2 Gy) at developmental Days 14 and 56, and samples were analyzed at 2 h, 6 h, 24 h, and 14 d post-irradiation. (A-D) Proliferation was analyzed by immunostainings for phosphorylated histone H3 (PH3) and phosphorylated Vimentin (pVim). (E-H) Cell death was accessed by immunostaining for the early apoptotic marker cleaved caspase-3 (CC3), and the late apoptotic marker TUNEL. n = 4. Image analysis was performed using the ImageJ software. One- way ANOVA test followed by Tukey’s test for multiple comparisons was used when comparing 3 conditions, *p<0.01 **p< 0.009; ***p< 0.0007; ****p< 0.0001. Student’s t-test was used when comparing 2 conditions. Scale bar = 50 µm.

When DNA damage remains unrepaired following cell cycle arrest, p53 activation can lead to apoptosis, a protective mechanism that eliminates irreparable cells, thus preventing genomic instability (57). To further investigate the mechanisms by which IR-induced DNA damage affects forebrain organoid growth, apoptosis was evaluated as a possible outcome of the activation of p53.

Immunostainings for the early and late apoptotic markers, cleaved caspase-3 (CC3) and TUNEL, respectively, were performed. At 6 h post-irradiation, apoptotic cells (CC3+ or TUNEL+) were observed at both developmental time points (Figure 3E-H). However, this was more pronounced in D14 organoids compared to D56 organoids (Figure 3E,F). The D14 organoids also presented a dose-dependent response (Figure 3G), which was not observed in D56 organoids, where significant cell death occurred only at the 2-Gy dose (Figure 3H). At 24 h, cells were no longer entering apoptosis (CC3+), although cells in final apoptotic stages were still present (TUNEL+) (Figure 3G,H). These findings reinforced the increased sensitivity of early stages of brain development to DNA damage compared to more developed stages.

Premature differentiation leading to an earlier generation of neurons instead of NPCs is another known mechanism underlying microcephaly (4). Recent studies have shown that exposure to DNA damage during early embryonic brain development in mice can also trigger premature differentiation (23). To investigate whether NPCs within human forebrain organoids undergo a premature switch from proliferative to differentiative cellular division following IR exposure we performed immunostainings for SOX2, PAX6, and DCX. Additionally, cells labeled with 5-bromo- 2’-deoxyuridine (BrdU) before exposure to IR were also stained. As organoids generated at developmental age D14 did not yet express PAX6, we accessed the expression of SOX2 that showed a reduction in the number of positive cells in a dose-dependent manner as early as 6 h following IR exposure (Figure 4A,C). At D56, the number of cells expressing PAX6 was also reduced depending on the radiation dose, however, the response was only apparent 24 h after irradiation (Figure 4A,C). To understand whether this reduction in the progenitor pool was due to premature differentiation, we then assessed DCX expression, a marker for immature neurons. We observed increasing fluorescence intensity of DCX in D14 organoids after 6 h (Figure 4B,D), and this effect could still be seen 14 days following irradiation exposure (Figure 4D). At D56, we could observe a slight increase in DCX but that was not statistically significant (Figure 4B,D), perhaps due to the reduced numbers of neural rosettes evaluated for this specific experiment. To confirm these findings, replicating cells (BrdU+) were investigated for their co-expression of DCX. Interestingly, organoids exposed to IR presented increased amounts of BrdU+DCX+ expressing cells at both developmental stages (Figure 4E,F). In D14 organoids, increasing numbers of BrdU+DCX+ cells were observed 24 h after IR exposure, with a more pronounced increase at a 0.5-Gy dose (Figure 4E,F). In contrast, in D56 organoids, this increase was more pronounced 14 days following IR exposure, in a dose-dependent manner (Figure 4E,F). We believe that the increased amount of cell death observed in D14 organoids exposed to a 2-Gy dose, might have contributed to the reduced number of BrdU+DCX+ cells observed for this dose, which was not the case for D56. Overall, these findings indicate that actively replicating cells that incorporated BrdU at the time of IR exposure, were increasingly committed to becoming neurons upon further evaluation.

**Figure 4.**
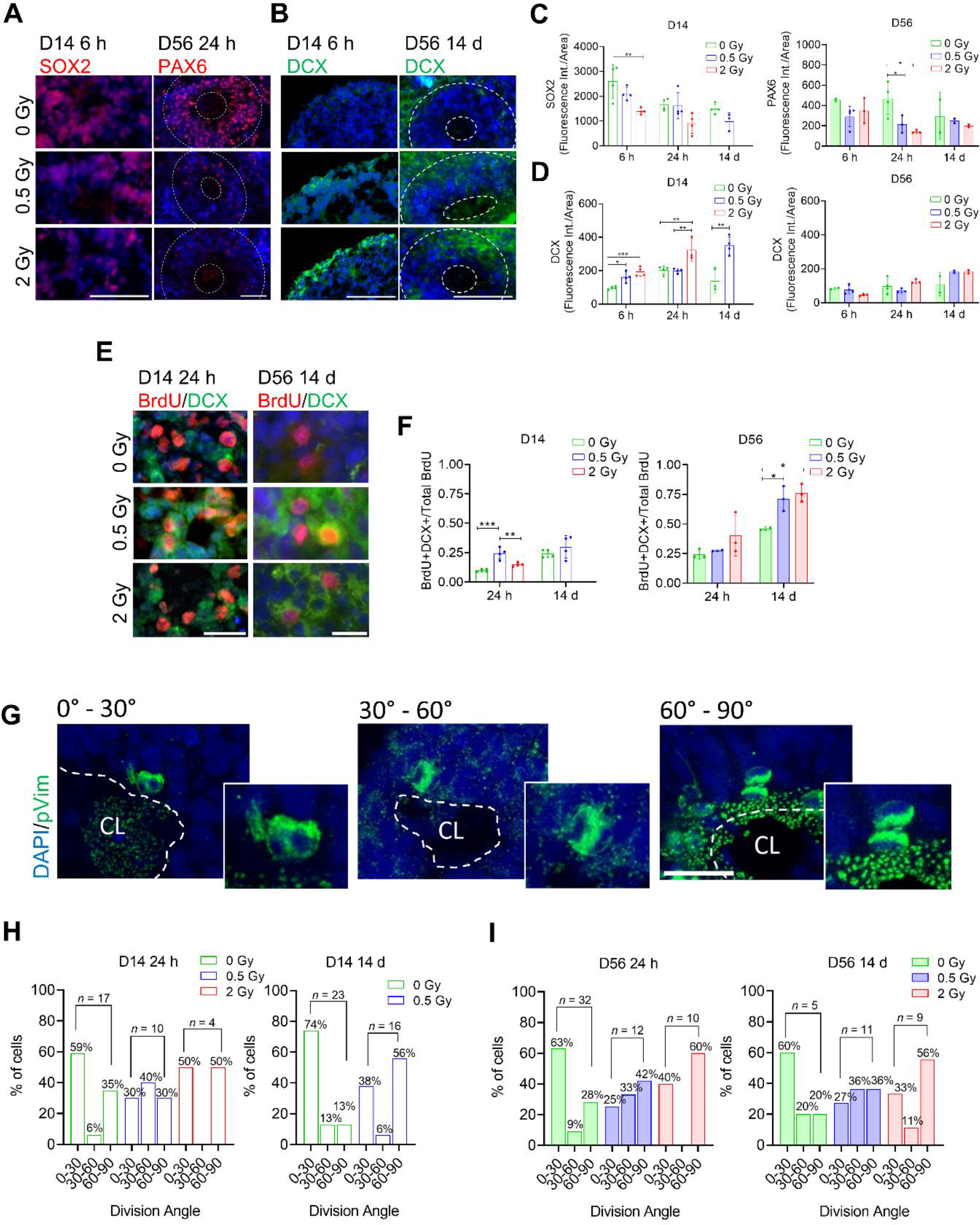
Ionizing radiation-induced DNA damage leads to premature differentiation in human forebrain organoids. Forebrain organoids were exposed to ionizing radiation (Sham-irradiated x 0.5 Gy x 2 Gy) at developmental Days 14 and 56, and samples were analyzed at 2 h, 6 h, 24 h, and 14 d post-irradiation. (A-D) Neural progenitor cells were accessed by immunostainings for SOX2 in D14 organoids and PAX6 in D56 organoids, while immature neurons were accessed by immunostainings for DCX at both developmental stages. Scale bar SOX2, PAX6 = 50µm, DCX = 100 µm. (E, F) Actively proliferating cells were labeled with BrdU 2 h before IR exposure. Following irradiation, immunostainings for BrdU and DCX were used to identify actively proliferating cells committed to becoming neurons. Scale bar = 50 µm. (G) Mitotic pVim+ cells were scored based on their division angle, with the central lumen (CL) of the neural rosette as the reference point: 0-30° is indicative of symmetric cellular division, while 30-60° and 60-90° are indicative of asymmetric cellular division. n = 4. Image analysis was performed using the ImageJ software. One-way ANOVA test followed by Tukey’s test for multiple comparisons was used when comparing 3 conditions, *p<0.04 **p< 0.008; ***p< 0.0008; ****p< 0.0001. Student’s t-test was used when comparing 2 conditions, **p< 0.005.

During neurogenesis, a balance between proliferative symmetric and differentiative asymmetric cell divisions ensures the generation of an adequate number of NPCs and, subsequently, neurons. When this balance is disrupted and cells switch from symmetric to asymmetric divisions earlier than expected, premature differentiation takes place. Here, we investigated cell division symmetry by scoring mitotic pVim+ cells within neural rosettes into three division angle categories: 0-30°, 30-60°, and 60-90° (Figure 4G). This allowed for the identification of the type of division that cells underwent following IR exposure. As expected, sham-irradiated D14 and D56 organoids displayed more cells undergoing symmetric cellular divisions (0-30°) than cells undergoing asymmetric divisions (30-60° or 60°-90°) (Figure 4H,I). However, following irradiation, we observed a dose-dependent shift in the division angle, with increased numbers of cells dividing asymmetrically at both developmental time points (Figure 4H,I). This effect was more pronounced 14 d after IR exposure. These findings confirm premature differentiation of NPCs following DNA damage exposure in humans.

### Transcriptional profiling confirms the activation of a dose- and developmental timing-dependent, p53-regulated DNA damage response in irradiated organoids

To further investigate molecular mechanisms responsible for the observed changes after irradiation of organoids, RNA-seq was performed on organoids irradiated at D14 or D56 and processed for RNA extraction after 6 h, 24 h and 14 days. These time points reflect both the early and late response to radiation. Because no effect of radiation could be observed via qRT-PCR on the expression of typical p53 target genes at 2 h after exposure (Figure S4A), this very early time point was not used for RNA-seq.

Principal component analysis indicated that the variation in gene expression profiles was mainly dependent on the developmental age of the organoids (Figure S4B-C) while the radiation dose contributed much less to the variation (Figure S4D). The overall effect of radiation on gene expression was most pronounced at 14 days after irradiation of D14 organoids irradiated with 2 Gy (Figure S4D), which corresponded to those organoids most affected in terms of size (Figure 2E).

Differential expression analysis (Table S2) indicated a clear dose-dependent response in gene expression at all time points. Both the number of DEGs (FDR <0.05) (Figure 5A) as well as their fold change (data not shown) were generally increased with the higher dose. A further indication of a dose response could be deduced from the overall low numbers of genes that were significantly differentially expressed after irradiation with 0.5 Gy, but not 2 Gy, while many more genes were either regulated at both doses or at 2 Gy only (Figure S5A,D; Table S2). Overlap analysis between different conditions using RedRibbon (Figure 5B) and categorical DEG list comparisons (Figure 5C,D) indicated that very significant overlaps existed between both up-and downregulated genes after 0.5 Gy and 2 Gy, especially at the earlier time points after exposure (6 h and 24 h). Also, a high overlap existed between upregulated DEGs after 6 h and 24 h of organoids irradiated at D14 and D56. Overall, the overlaps in different experimental conditions between upregulated genes were more significant than the overlaps between downregulated genes (Figure 5C,D).

**Figure 5.**
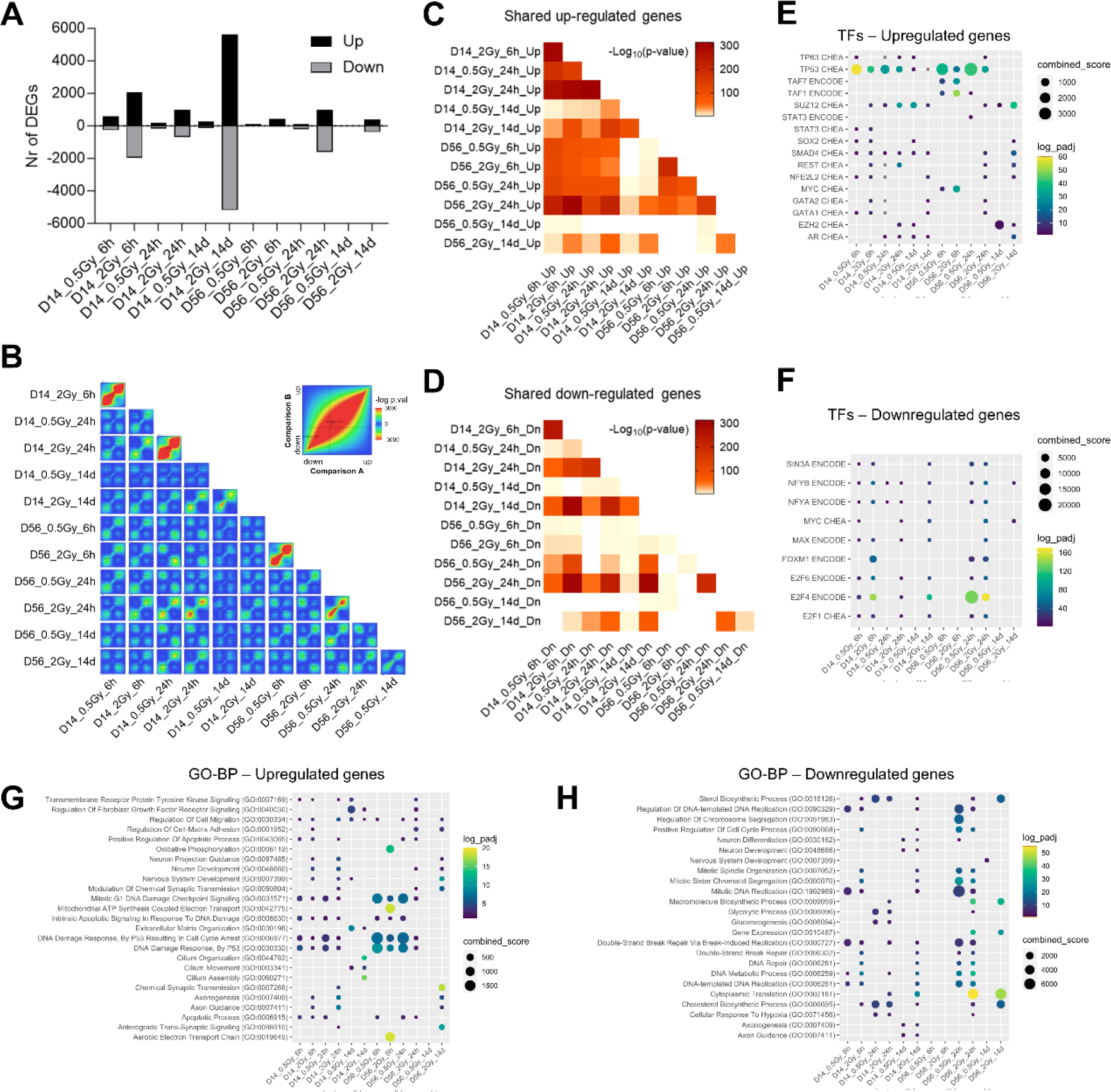
Dose- and time-dependent changes in gene expression after irradiation of organoids. (A) Number of significantly (FDR < 0.05) up- (black bars) and downregulated (gray bars) differentially expressed genes (DEGs) in irradiated versus sham-irradiated organoids at each experimental time point. (B) RedRibbon hypergeometric overlap maps for the different experimental comparisons. For each comparison, the indicated condition was compared to its respective 0-Gy control condition. On each map, the extent of shared downregulated genes is displayed in the bottom left corner, whereas shared upregulated genes are displayed in the top right corners. Inset indicates perfect overlap. (C, D) Statistical hypergeometric overlap between lists of up- (C) and downregulated (D) DEGs among the different experimental comparisons. (E, F) Enrichment analysis of predicted transcriptional regulators of up- (E) and downregulated (F) DEGs.(G, H) Enrichment analysis of Biological Processes of up- (G) and downregulated (H) DEGs. See also Figure S4-S9.

Upregulated genes at both D14 and D56 were mostly predicted to be regulated by p53, especially those genes that were upregulated at the early time points (Figures 5E and S5B). Genes upregulated at the later time points in both D14 and D56 organoids were enriched in targets of REST and the PRC2 complex members SUZ12 and EZH2, suggesting an epigenetic component in their regulation. Downregulated genes, on the other hand, were predicted to be regulated mostly by members of the E2F family, especially by the transcriptional repressor E2F4 (Figure 5F). Other predicted regulators of downregulated genes were NFYA/B, MYC and MAX. Altogether, radiation-induced genes were regulated by TFs that also regulate developmentally induced genes (Figure S1C), while radiation-repressed genes were regulated by TFs that regulate developmentally repressed genes (Figure S1E).

In terms of pathways affected by radiation, early upregulated genes after 0.5 Gy and 2 Gy were involved in DNA damage response, cell cycle arrest and apoptosis (Figure 5G). This corresponded well with observations from immunostainings (Figures 2,3) and the activation of p53. However, early gene upregulation that was more specific to the 2-Gy dose was related to neuronal differentiation pathways (e.g. neuron development, nervous system development and axon guidance) (Figures 5G and S5B). This effect was also more pronounced in D14 compared to D56 organoids. Activation of these genes may be related to the observed premature neuronal differentiation. From the numbers of DEGs at each experimental condition and their overlaps between different conditions, we could also deduce that the transcriptional response to radiation in D56 organoids was both delayed and attenuated as compared to that in D14 organoids. Indeed, while in D14 there was a very strong response after 6 h (859 DEGs – 0.5 Gy and 4047 DEGs – 2 Gy), which diminished after 24 h (360 DEGs – 0.5 Gy and 1641 DEGs – 2 Gy), the opposite was seen in D56 organoids with a much smaller response after 6 h (128 DEGs – 0.5 Gy and 490 DEGs) compared to that after 24 h (316 DEGs – 0.5 Gy and 2580 DEGs – 2 Gy) (Figures 5A and S5A,D). Pathways that were enriched after both 6 h and 24 h in D14 organoids, were only enriched after 24 h in D56 organoids (Figure S5B,C), again indicating a delayed response.

In addition to differential expression analysis, we also inferred gene regulatory networks using LemonTree, an ensemble method that first groups genes into coexpression modules and then assigns regulatory TFs to these modules. The clustering indicated nine modules consisting of genes that showed an obvious dose-dependent increase in expression (Figure S6A,B). In the majority of cases the induction of these genes was most pronounced at 6 h after exposure of D14 organoids, with either a reduced (modules 32, 34, 55, 61, 93, 162) or similar (modules 69, 70, 145) transcriptional response after 24 h. In D56 organoids, the effect on expression of these genes was reduced and sometimes absent, again indicating an attenuated response. In terms of enrichment, these upregulated genes were highly enriched as predicted targets of p53 and involved in typical p53-dependent DDR processes and apoptosis (Figure S6C,D).

Early (6h, 24 h) downregulated genes in D14 organoids, on the other hand, were involved in pathways related to DNA replication, cell division and DNA repair (Figures 5H, S5E,F and S7). This is consistent with their regulation by the transcriptional repressor E2F4 (Figure 5F). Other enriched pathways among downregulated genes, primarily after 24 h, were related to metabolism (sterol biosynthesis, gluconeogenesis, fatty acid biosynthesis) and these were regulated by NFY family TFs (Figure 5H and S8). The dose-dependent downregulation of these metabolic pathways was a surprising result. It is unclear how this may have influenced the growth of irradiated organoids, however, it is well established that disruptions of metabolic pathways like cholesterol biosynthesis and glycolysis can result in neurodevelopmental defects including microcephaly (58).

While irradiation of D14 organoids with 0.5 Gy induced persistent changes in gene expression (256 genes up, 130 genes down), almost no long-term changes (3 genes up, 18 genes down) were observed in D56 organoids irradiated with the same dose. This is another indication of the relative insensitivity of older organoids to a moderate radiation dose which resulted in levels of DNA damage that could be efficiently repaired and therefore did not induce substantial levels of apoptosis (Figure 3H).

Among the genes that were upregulated 14 days after exposure of D14 organoids were also many genes related to primary cilia assembly and motility (Figure 5G), including members of the cilia and flagella associated protein (CFAP), sperma associated antigen (SPAG) and dynein arm (DNAAF, DNAH, DNAI and DNAL) families (Figure S9A). In the developing brain, the ChP epithelium consists of multiciliated cells (59), responsible for circulation of cerebrospinal fluid (60). Thus, we examined expression of markers of the ChP and found a remarkable increase of their expression at 14 days after exposure of D14 organoids (Figure S9B). This coincided with changes in genes belonging to the WNT/FZD pathway (Figure S9C) which regulate the epithelial fate of the ChP (61). This suggested that ChP cells were spared from radiation-induced apoptosis, consistent with their resistance to traumatic injury (61).

### Irradiation of forebrain organoids reduces expression of MCPH genes

Enrichment analysis of disease-associated genes from DisGeNET and Jenssen Disease databases, showed that genes associated with neurologic diseases and syndromes characterized by reduced brain size, like Fanconi Anemia, Seckel syndrome and MCPH were highly enriched among downregulated genes, both in D14 and D56 organoids (Figure 6A,B). Thus, we hypothesized that downregulation of these genes may be a contributing factor to the growth deficit we observed in irradiated organoids.

**Figure 6.**
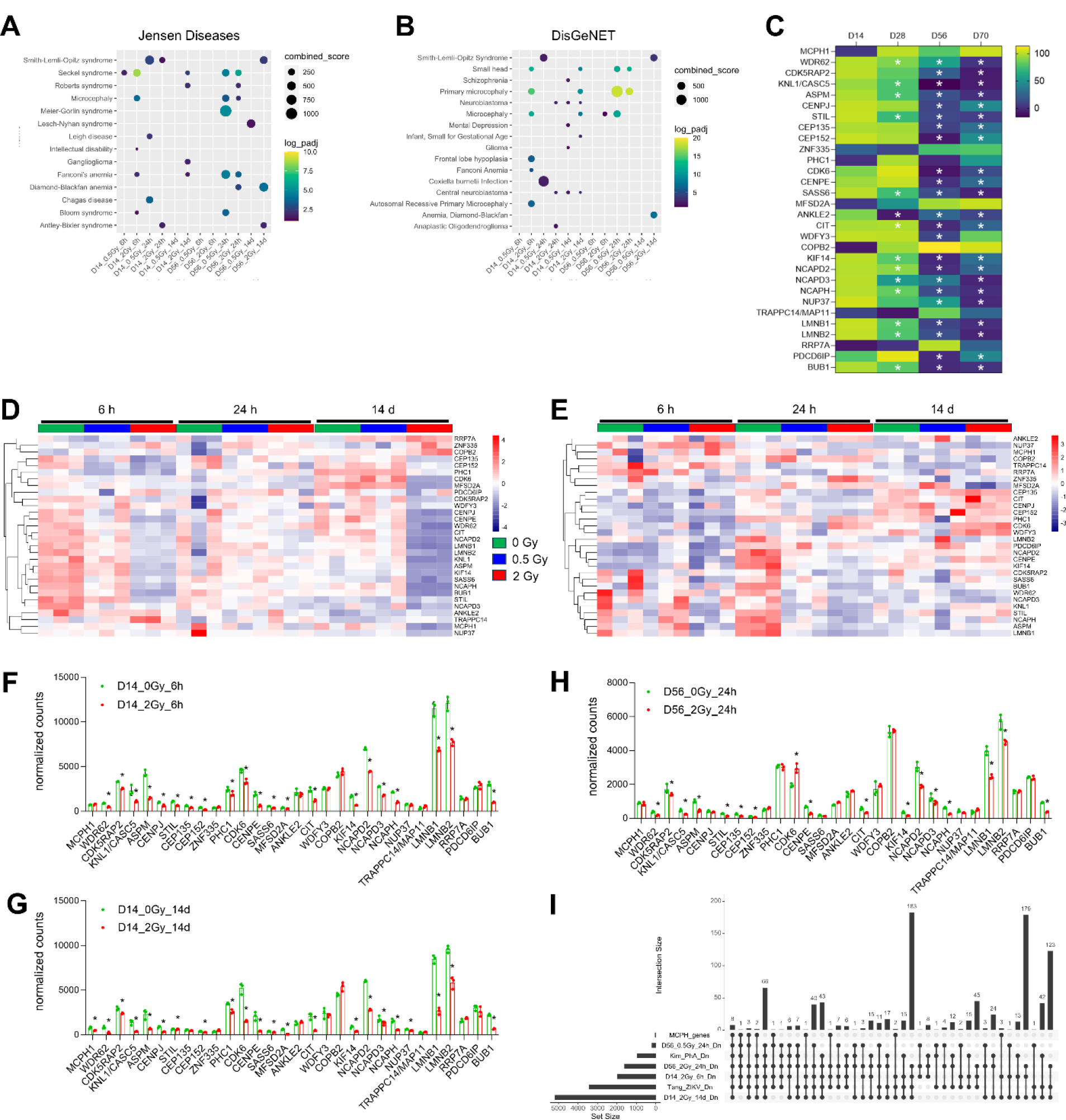
Microcephaly (MCPH) genes are coordinately downregulated during organoid development and after irradiation. (A, B) Enriched neurological diseases and syndromes associated with reduced brain size among radiation-repressed genes from Jensen_Diseases (A) and DisGeNET (B) databases. (C) Heatmap of MCPH gene expression at different stages of organoid development. *denotes significant downregulation (FDR <0.05) in comparison to D14. (D, E) Heatmaps of MCPH gene expression after irradiation of D14 (D) and D56 (E) organoids. (E-H). Normalized RNA-seq counts in 0-Gy (green bars) and 2-Gy (red bars) irradiated organoids. *denotes significant differential expression (FDR <0.05) in comparison to 0 Gy. (I) UpSet plot indicating overlaps between MCPH genes, radiation-repressed genes in organoids at different time points, repressed genes in human NPCs after infection with Zika virus (From (64)), and repressed genes after treatment of human brain organoids with phenylalanine (From (65)) Intersections were ranked according to degree of overlap. Note that not all overlapping intersections are shown. See also Figure S10, S11.

Therefore, we analyzed gene expression profiles of the 30 known MCPH genes (6,8,9) in our organoids, both during their development and after irradiation. We found that the majority of MCPH-associated genes were significantly downregulated during organoid maturation, with the biggest changes occurring between D28 and D56 (Figure 6C). This corresponds to their preferential expression in radial glia (Figure S10) (62), which are the NPCs of the cortical ventricular zone.

In response to radiation, many of the MCPH genes showed highly similar expression profiles (Figure 6D-H): in D14 organoids, 21 MCPH genes were downregulated 6 h after a 2-Gy dose (Figure 6D,F) and 22 MCPH genes were highly reduced after 14 days (Figure 6D,G). In D56 irradiated organoids, however, 16 MCPH genes were dose-dependently reduced after 24 h, but showed normal expression after 14 days (Figure 6E,H). Fourteen of these genes were commonly downregulated at all these experimental conditions. Moreover, the same genes were also downregulated in human NPCs that were infected with the microcephaly-inducing Zika virus (Figure 6I) (63,64). Likewise, treatment of brain organoids with high doses of phenylalanine caused a growth reduction which coincided with activation of the p53 pathway, apoptosis and decreased expression of several MCPH genes (65), all of which were also affected by irradiation (Figure 6I). Thus, irradiation induced a concerted repression of MCPH genes that could have contributed to the observed growth deficits.

One possible explanation for the long-term effect of radiation on MCPH gene expression in D14 organoids could be the apparently changed cellular composition of these organoids, with an enrichment in ChP and CR cells (Figure S9). However, based on single-cell RNA-seq data of MCPH genes in different cell types of the developing mouse brain, including ChP and CR, (66) (66), this seems very unlikely. Several of the radiation-repressed MCPH genes (*Cdk5rap2*, *Stil*, *Cep135*, *Cep152*, *Sass6*, *Cit*, *Ncaph*) actually show highest expression levels in ChP compared to other cell types (Figure S11). So, on the basis of cellular composition one would expect an upregulation of these genes, and therefore the downregulation we have observed here might even be an underestimation.

### Radiation-induced repression of microcephaly genes is p53-E2F4/DREAM-dependent and occurs in human, but not mouse NPCs and embryonic brain

In our previous studies on radiation-induced microcephaly in mice (23,32) we had never noticed a reduced expression of microcephaly genes that could contribute to the microcephalic phenotype. In fact, in those studies we generally found very few downregulated genes in comparison to those that were upregulated. Therefore, we re-analyzed those datasets, with a focus on MCPH genes. This showed that indeed, much fewer MCPH genes were significantly downregulated in irradiated mouse embryos and NPCs compared to human organoids, with little to no overlap between different experiments (Figure S12A-C).

However, in our previous studies in mice, the highest radiation dose used was 1 Gy, which was sufficient to induce massive apoptosis and persistent brain size reduction (22,23). To verify that the lower dose could be the reason for the observed discrepancy in gene expression profiles between mouse embryos/NPCs and human organoids, we irradiated human and mouse NPCs with a dose of 1 Gy and analyzed gene expression of selected p53 targets and MCPH genes at 6 and 24 h following exposure using qRT-PCR. As MCPH genes we chose *ASPM*, *WDR62*, *CIT* and *KNL1*. Biallelic mutations in *ASPM* and *WDR62* are the most common causes of MCPH (68.6% and 14.1%, respectively) (67), and *ASPM*/*Aspm* expression was found to be reduced by radiation in human fibroblasts and fetal mouse brain and neurospheres (68). *CIT* and *KNL1* are both involved in mitotic spindle organization and are located in the spindle poles and the kinetochore, respectively. Genetic mouse models of *Cit* and *Knl1* display microcephaly as a consequence of p53-mediated apoptosis of neural progenitors (69,70). During early neurogenesis, fetal brains of these mice display gene expression profiles that are very similar to those of irradiated mouse fetuses (70–72).

The p53 targets *CDKN1A* and *EDA2R* were induced ∼6-fold and ∼4-fold, respectively, in human NPCs (Figure 7A). In mouse NPCs they were only induced 3-fold and 2.5-fold (Figure 7B). In contrast, in human NPCs the selected MCPH genes showed overall a moderately reduced expression after 6 h (except for *KNL1*), which further reduced to ∼25% after 24 h (Figure 7C). In mouse NPCs however, no change in expression was observed, except for a transient reduction of *Wdr62* which returned to normal levels after 24 h (Figure 7D). Thus, radiation-induced repression of the MCPH genes was specific to human NPCs.

**Figure 7.**
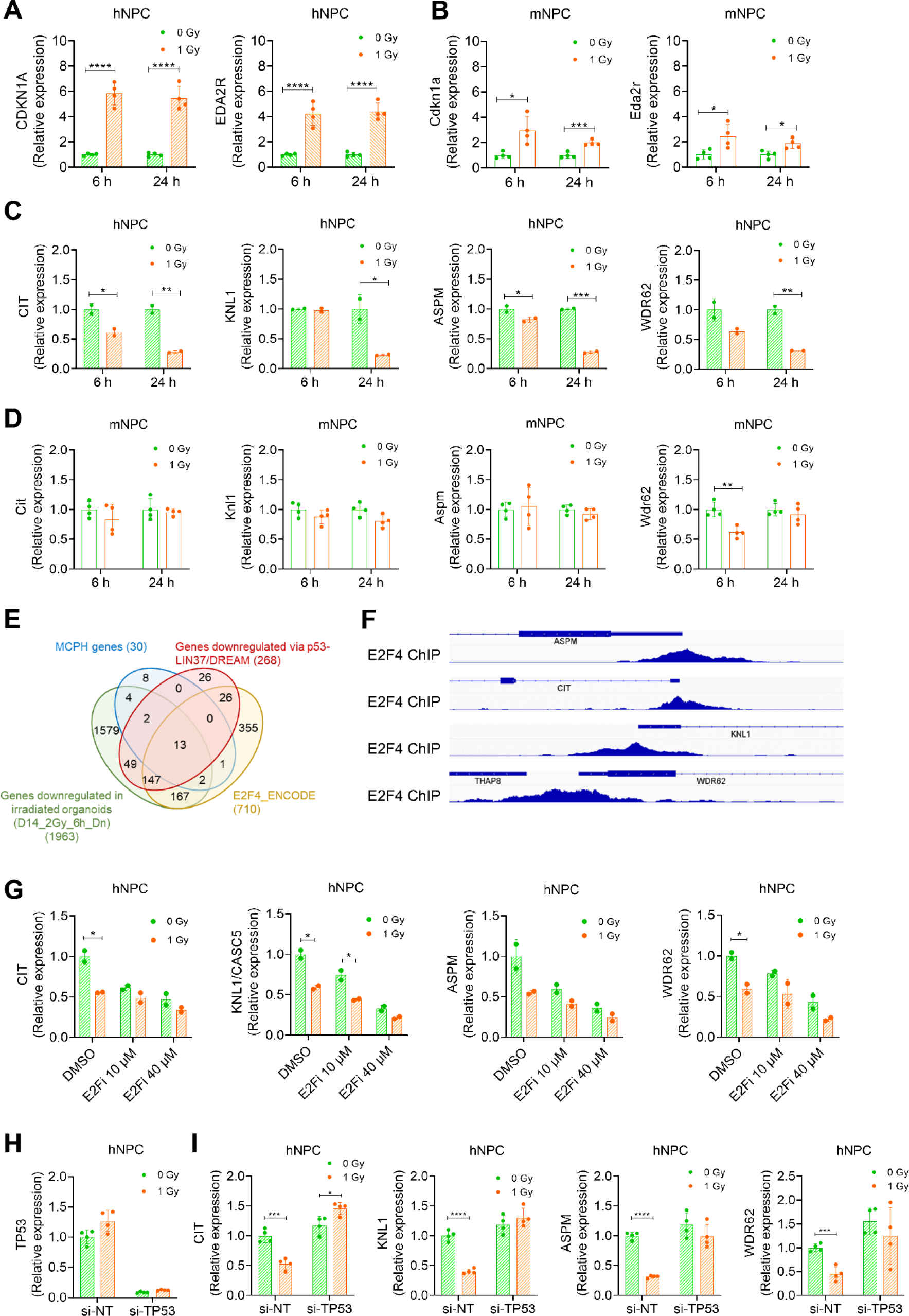
Radiation-induced repression of MCPH genes is human-specific and dependent on p53-E2F4/DREAM. (A, B) Radiation-induced activation of the p53 target genes CDKN1A and EDA2R in human NPCs (A) and Cdkn1a and Eda2r in mouse NPCs (B) (n = 4). (C, D) Selected MCPH genes are repressed after irradiation in human NPCs (C) (n = 2), but not in mouse NPCs (D) (n = 4). (E) Overlaps between early repressed genes in D14 organoids, MCPH genes, targets of p53-LIN37/DREAM and E2F4 targets. (F) E2F4 ChIP-seq tracks at promoters of selected MCPH genes retrieved from ChIP-Atlas (www.chip-atlas.org). (G) MCPH gene expression is reduced after treatment of human NPCs with an E2F inhibitor, which has no effect on their repression after IR (n = 2). (H) Silencing of TP53 in human NPCs almost completely ablates its mRNA expression. (I) Silencing of TP53 prevents radiation-induced repression of MCPH genes (n = 4). Student’s t-test was used,**** p<0.0001, ***p< 0.0008, **p< 0.007, *p< 0.04. See also Figure S12, S13.

Previous studies have shown that activation of p53 (e.g. via DNA damage induction) leads to transcriptional repression of indirect p53 targets via the DREAM complex which is composed of the MuvB core complex, E2F4-5/DP, and p130 or p107. E2F4 and E2F5 are transcriptional repressors that bind to E2F sites, while p130 and p107 are activated by p21/CDKN1A in a p53- dependent way. Thus, DREAM, via the E2F4 or E2F5 subunits has been shown to coordinately repress many genes in a p53-dependent manner, especially those involved in cell cycle regulation but also DNA repair or Fanconi Anemia, as well as epigenetic regulators like EZH2 and SUZ12 (18). Furthermore, a recent study found that p53-dependent repressed genes were either regulated via LIN37/DREAM (268 genes) or not (415 genes) (73). Interestingly, we noticed that 15 out of 30 MCPH genes are among the 268 LIN37/DREAM-dependent genes while no MCPH gene belongs to the list of genes downregulated independent of LIN37/DREAM. Moreover, all 15 LIN37/DREAM-dependent MCPH genes were also downregulated in our irradiated forebrain organoids, while of the 15 LIN37/DREAM-independent MCPH genes only 6 were downregulated in our study (Figure S12D,E). Likewise, 16 MCPH genes are known targets of E2F4, the main regulator of gene repression in irradiated organoids (Figure S12F,G), all of which were among the downregulated ones and E2Fs, including E2F4, were identified by motif enrichment analysis of gene modules containing MCPH genes (Figure S12H). Finally, enrichment analysis using ChIP- Atlas (www.chip-atlas.org) (74) indicated that E2F4 was the most highly enriched TF for MCPH genes in neural cells (Q-value = 10E-9.7; Fold Enrichment 12.2) and E2F4 binds the promoters of *CIT*, *KNL1*, *ASPM* and *WDR62* (Figure 7F). Altogether, this suggested that the coordinated repression of MCPH genes in irradiated organoids occurred in a p53-E2F4/DREAM-dependent manner.

To investigate this, we treated human NPCs with the non-selective E2F inhibitor (E2Fi) HLM006474 before (sham-)irradiation and performed qRT-PCR for the selected genes. This showed an E2Fi dose-dependent downregulation of MCPH genes, although E2Fi did not rescue their radiation-induced repression (Figure 7G). Therefore, we knocked down *TP53* in human NPCs (Figure 7H) 24 h before irradiation. This led to an ablation of radiation-induced MCPH gene repression (Figure 7I). Altogether, these results indicate that coordinated MCPH gene repression in human organoids and NPCs is regulated by p53-E2F4/DREAM.

Concerning the apparent differential regulation of these genes in human versus mouse NPCs, it should be noted that p53-DREAM-dependent gene repression but not direct p53-dependent gene activation, was highly conserved between humans and mice in response to different treatments that result in p53 activation (75). A curated database of human and mouse p53-regulated genes exists (www.targetgenereg.org; (76)) which contains data about p53-dependent gene activation or repression as well as chromatin immunoprecipitation results of transcriptional activators (e.g. p53) or repressors (e.g. E2F4, DREAM). Therefore, we interrogated the TargetGeneRegulation database for p53-dependent expression and DREAM- (human) or E2F4-dependent (mouse) regulation of MCPH genes (75). This indicated that both human and mouse MCPH genes show an inverse correlation between P53 Expression Score and DREAM- or E2F4-binding Score, respectively (Figure S12I). However, while human radiation-repressed genes were highly enriched among genes with a high DREAM Score and low P53 Expression Score, this was not the case for the few mouse radiation-repressed MCPH genes (Figure S12I). Surprisingly, when we treated mouse NPCs with E2Fi, we also found a significant downregulation of the selected MCPH genes, which was prevented in NPCs derived from conditional *Trp53* knockout mice (Figure S13A). As in the previous experiments, irradiation had little to no effect on MCPH gene expression in mouse NPCs. Thus, inhibition of E2F influences MCPH gene expression in mouse NPCs in a p53-dependent manner, as it does in human NPCs, although this is not the case in response to radiation exposure. This contrasts previous findings in mouse embryonic fibroblasts (MEFs), where expression of the MCPH genes *Knl1*/*Casc5*, *Wdr62*, and *Ncaph* were significantly downregulated after treatment with the p53 activator Nutlin in a p53-DREAM-dependent manner (77). Therefore, we performed irradiation experiments in MEFs in the presence or absence of E2Fi. This showed that E2Fi treatment severely reduced MCPH gene expression in wild-type MEFs, although again, irradiation had little to no effect on MCPH gene expression (Figure S13B).

Altogether, this suggests that the difference in MCPH gene regulation between human organoids and mouse embryonic brains, and human and mouse NPCs may be context dependent, and specific to the type of DNA damage and the subsequent response inflicted by acute exposure to ionizing radiation.

## 3. Discussion

In this study, hESC-derived forebrain organoids were used as a model system to deepen our understanding of DNA damage-associated microcephaly in humans. Regarding underlying processes, it is currently known that the hyperactivation of p53 leading to exacerbate apoptosis and NPC pool depletion is the main mechanism accounting for endogenous and exogenous DNA-damage induced microcephaly in mice (23,33). Here, we used acute exposure to ionizing radiation to induce DNA damage and demonstrated that this resulted in a reduction in organoid size, which was shown to be dose- and developmental-time dependent. Besides a classical DDR and the induction of premature neuronal differentiation, we unexpectedly found a coordinated downregulation of many MCPH genes in irradiated organoids, which was p53-E2F4/DREAM dependent and specific to human cells.

Human forebrain organoids are recognized as a powerful tool to model different neurodevelopmental disorders (78–80), given their ability to recapitulate many characteristics of early cortical development. However, differences between normal brain development and organoid development have been reported (81–83), and obviously, organoids lack many of the specific characteristics of the in vivo environment. Nevertheless, using immunostainings and RNA-seq we could validate that the organoids used in our study shared many features with organoid cultures from other studies in which they were used to investigate aspects of human neurodevelopment.

So far, the underlying mechanisms leading to radiation-induced microcephaly have only been investigated in animal models (21,22,31,32,39). Lately, efforts have been made to understand some of the mechanisms based on the principle of Adverse Outcome Pathways, which hinted at some affected pathways (84,85), although without experimental findings from human studies the picture would never be complete. A few studies have previously used the organoid model to investigate radiation effects, however, none of these studies were performed in the context of radiation-induced microcephaly and mostly focused on DNA damage repair kinetics (40,41,54). Oyefeso *et al.,* also performed RNA-seq at 48 h post-IR. The findings of these studies, were overall in line with some of our own observations, namely a dose- and time-dependent efficiency of DNA repair, and the activation of genes related to neuronal pathways while cell cycle genes were reduced (40).

### Irradiation of organoids results in the activation of the DDR, which is p53- and E2F4- dependent, and culminates in reduced organoid growth

Following IR exposure, impaired neural progenitor proliferation and neurogenesis was evident by a dose- and developmental-time-dependent reduction in organoid size, with D14 organoids being highly affected after exposure to a 2-Gy dose. Organoids irradiated with 0.5 Gy could recover their growth capacity over time, but could not reach the size of sham-irradiated controls, for both developmental time points. Indeed, it has been previously described that higher doses of IR exposure increase the risk of neuronal defects, and that the early developing human brain is extremely sensitive to DNA damage exposure compared to more advanced stages, increasing the susceptibility to microcephaly (11,86). Additionally, structural alterations of neural rosettes were observed, with irritated D14 organoids displaying enlargement and thickening of these structures. We believe this may be related to the loss of cellular architecture, as we observed atypical rosette morphology, accompanied by an extensive cell death in D14 organoids. In contrast, irradiated D56 organoids exhibited smaller rosettes, both in terms of perimeter and thickness. We attribute this effect to reduced proliferation and premature differentiation of NPCs, as no morphological changes within rosettes were apparent at this stage. In line with our findings, a recently published study modeling a neurodevelopmental disorder in forebrain organoids demonstrated that a reduction in neural rosette size and thickness is associated with deficient proliferation (87).

A typical dose-dependent DDR, regulated primarily via p53 (upregulated genes) and E2F4 (downregulated genes) was observed after irradiation. This was characterized by p53 activation, DNA repair, cell cycle arrest, and apoptosis, on the basis of our immunostaining experiments. Organoids at earlier developmental stages appeared to be more proficient in repairing DNA damage, as γH2AX and 53BP1 foci were resolved faster compared to mid-development stage organoids (Figure 2A,,B). Indeed, it has been previously described that SOX2+ NPCs are more efficient in repairing damage to the DNA when compared to mature CTIP2+ neurons (54). However, we believe that some of the highly IR-sensitive progenitors within D14 organoids, having accumulated too much DNA damage, had already undergone apoptosis by 6 h, reducing the number of apparent foci in our analysis. Indeed, D14 organoids displayed an extensive number of CC3+ and TUNEL+ cells, especially after a 2-Gy dose, while D56 organoids displayed only mild apoptosis. Thus, a combination of both an increased DNA repair capacity and increased apoptosis contributed to the apparent faster resolution of DNA damage foci in D14 organoids. Along with this, the response to radiation exposure was delayed and attenuated in D56 organoids compared to D14 organoids. This was supported by the prolonged, but less extensive cell cycle arrest in D56 organoids, the reduced number of apoptotic cells (Figure 3) and the transcriptomic analysis. An attenuated DDR in older organoids, is in line with the higher DNA damage sensitivity of the brain during early neurogenesis and could be due to the decreased threshold of early neural progenitors to induce apoptosis (88), or differences in their cell cycle dynamics, i.e. shorter G1 phase, which renders them more susceptible to genotoxic stress (89).

Another feature that is often seen in models of microcephaly is premature neuronal differentiation (4) which also leads to a depletion of proliferating progenitors. Here, we performed immunostainings for neural progenitor markers (SOX2, PAX6), early post-mitotic neuron markers (DCX) and we followed-up on the fate of cells that were proliferating at the time of exposure via BrdU. Altogether, this showed, in agreement with our previous findings in mice (23), that irradiation led to a reduction in NPCs (possibly partly due to apoptosis), an increase in DCX+ neurons and premature cell cycle exit, both at D14 and D56 (Figure 4). These findings were supported by our transcriptomic analysis demonstrating a general increase in expression of genes related to neuronal differentiation processes, many of which were predicted to be regulated by TFs like REST and the PRC2 complex members SUZ12 and EZH2, which also regulate developmental gene expression. Furthermore, we found that the mitotic spindle orientation of dividing cells after IR had shifted to more asymmetric divisions (Figure 4H,I), and thus to premature differentiation (4).

Another interesting observation was the apparent enrichment of ChP cells in irradiated D14 organoids, which was evidenced by the high increase of ChP marker genes at 14 days following IR. This suggests that ChP cells are very resistant to DNA damage-induced apoptosis. A similar increase in ChP was seen in a recent pre-print article (90) in which D20 and D80 organoids were irradiated and showed an increased number of organoids with cavities (most likely corresponding to CSF-filled ventricles), especially after irradiation of D20 organoids. In our study, we did not specifically observe this. This could be due to the fact that our follow-up time was shorter, and the D14 organoids were likely much more prone to inducing apoptosis. In the study of Durante et al., changes in expression of some WNT genes were observed, which we also found here. Exactly why ChP cells would be so resistant to DNA damage is currently unknown. However, in one of our previous studies in mouse embryos we did see that ChP precursor cells were specifically protected from undergoing apoptosis after IR (Figure 2l in (23)).

### Human-specific repression of MCPH genes in irradiated organoids

An unexpected result from our study was the finding that irradiation of organoids resulted in the coordinated repression of a high number of MCPH genes. To date, 30 MCPH genes have been identified (6), and of these, 21 were downregulated in at least one condition, while 14 were downregulated in at least three experimental conditions. Importantly, a similar phenomenon was observed in human NPCs after infection with the Zika virus strain that caused microcephaly. This caused a proliferation defect in the cells and a global dysregulation of cell cycle-related genes (64), including many MCPH genes among which all but one were also dysregulated in our study. Although both radiation and Zika virus infection induce p53 signaling, it is unclear what specifically links the transcriptional responses to radiation and Zika virus, although we have previously shown that also among upregulated genes a high concordance exists between both environmental stresses (71).

Repression of genes in response to genotoxic insults, DNA damage, or chemotherapeutic drugs that affect p53 activity has been shown to be indirectly dependent upon p53 (91) via the activation of p21/CDKN1A. This leads to hypophosphorylation of the retinoblastoma protein RB and recruitment of p130 and E2F4, crucial members of the DREAM complex, to the promoters of cell cycle genes resulting in their transcriptional downregulation (92). Based on a recent study in HCT116 colon cancer cells (73) we found that 15 MCPH genes are negatively regulated by p53 in a LIN37/DREAM-dependent way. In contrast, among the genes that were repressed by p53 independently of LIN37/DREAM no MCPH gene was found, suggesting that MCPH gene repression primarily follows the p53-DREAM-dependent axis. Indeed, E2F4 targets and LIN37/DREAM-dependent genes were highly enriched among the genes repressed by radiation (this study), Zika virus infection (64), and phenylalanine treatment (51). We validated the involvement of E2F4/DREAM by pharmacological inhibition of E2Fs, resulting in downregulation of MCPH genes in NPCs (Figure 7G). This may seem a counter-intuitive result, since inhibition of a repressor would be expected to result in gene induction. However, HLM006474 is a non-selective inhibitor of E2Fs, which also inhibits the activator E2Fs, such as E2F1 (93). Moreover, in certain contexts, also E2F4 activates gene expression. This is the case for instance in embryonic stem cells where E2F4 activates transcription of cell cycle genes independent of RB. Thus, it is currently accepted that under conditions when RB is not activated, i.e. in the absence of p53/21 activation, E2F4 is mainly a transcriptional activator (94). This explains why E2F inhibition in non-irradiated NPCs resulted in the downregulation of MCPH genes. Also in glioblastoma stem-like cells E2F inhibition with HLM006474 resulted in downregulation of cell cycle genes (95). The role of p53 was established by knocking down *TP53*, which resulted in a rescue of the radiation-induced downregulation of MCPH genes (Figure 7I). Similar results were found in mouse NPCs, where E2Fi led to downregulation of MCPH genes while this was not the case in p53 knockout cells (Figure S13A). However, radiation-induced repression of MCPH genes was not observed (Figure S13A). Then why does radiation result in a different regulation of MCPH genes in human and mouse neuronal cells (human and mouse NPCs (this study), human forebrain organoids (this study), mouse embryonic brain (23,31) and primary neurons (31))? Especially, since a previous meta-analysis of the p53 gene regulatory network in human and mouse concluded that p53-dependent gene activation was less conserved than repression between the two species (75). One important factor may be the specific context of DNA damage and the following p53-E2F/DREAM activation dynamics induced by acute exposure to radiation. Perhaps this is one of the reasons why during the development of the TargetGeneReg database irradiation experiments were not included since they showed little overlap with the other datasets (96). An example of differences in context-dependent p53-DREAM regulation was seen in a recent study by Rakotopare *et al.* (77). Brip1 was identified as a p53-DREAM target gene, downregulated upon the differentiation of bone marrow cells and in ZIKV-infected NPCs, but not in irradiated hematopoietic stem cells. Interestingly, in our study BRIP1 was downregulated in irradiated organoids in all conditions in which we found MCPH gene downregulation, but also in irradiated mouse NPCs. In fact, in mouse NPCs radiation- repressed genes were also enriched in E2F4 targets, but these did not comprise the MCPH genes we found to be repressed in human organoids and NPCs.

Another explanation may be related to differences in cell cycle length between human and mouse neural progenitors (97). This may result in a different regulation of p53 activation dynamics and subsequent activation of p21/CDKN1A. In this respect, it is noteworthy that the amplitude of the activation of direct p53 targets, including CDKN1A, was reduced in mouse compared to human NPCs (Figure 7A, B). Whether that would also result in a difference in the recruitment of DREAM to gene promoters is currently unclear. Experiments comparing E2F4 recruitment to MCPH gene promoters after irradiation of human and mouse NPCs could potentially answer at least part of this question.

Therapeutic approaches to prevent (radiation-induced) microcephaly currently do not exist, although many studies in mice have indicated that it can be at least partly rescued by inhibition of p53 (8,9). However, targeting p53 is not without risk, given its role in many cellular processes. Therefore, MCPH gene targeting could perhaps be an additional avenue for possible treatment options. Interestingly, MCPH genes have been proposed as possible targets to treat glioblastoma as they control microtubule stability during cell division (98,99). One benefit of MCPH genes as targets is their restricted expression in NPCs, minimizing potential side effects.

## Limitations of the study

In this study, we used human forebrain organoids to evaluate the effects of radiation on their growth, aspects of the DNA damage response, cellular differentiation and molecular pathways. For this, we used an organoid culture protocol in which the embryoid bodies and subsequently the organoids are not embedded in an extracellular matrix, nor did we shake the cells as we observed that this led to their aggregation. This may have led to some degree of nutritional and oxygen deprivation, especially at the later stages of organoid development when they had reached a critical size. However, we do not believe that, if it happened, this would have influenced the main findings of our study, especially since we found similar results in monolayer NPC cultures. In any case, many different protocols for brain organoid cultures exist, each with their advantages and drawbacks (100,101). A recent study compared organoid cultures in the presence and absence of an exogenous extracellular matrix and they observed no major differences in morphology, cellular composition or gene expression profiles although organoids cultured in Matrigel did induce transcriptional pathways of eye development (102). Also, the fact that we used forebrain organoids rather than, for instance, dorsal-ventral assembloids or cerebellar organoids may have caused us to miss important aspects of the effects of radiation exposure on brain development. The reason for our choice was that so far, studies on radiation-induced (or DNA damage-associated) microcephaly in animal models have almost exclusively focused on cortical progenitors.

## Conclusion

In this study, we demonstrated that exposure of human forebrain organoids to acute doses of radiation induces a canonical DNA damage response culminating in apoptosis and premature differentiation of neural progenitors. This coincided with a downregulation of multiple genes linked to MCPH, altogether resulting in a reduced organoid growth, resembling microcephaly. Our findings revealed that the gene repression was indirectly driven by p53 via E2F4/DREAM, underscoring a unique mechanism that leads to neurodevelopmental defects in humans. Further research is necessary to understand how p53 activation dynamics induced by radiation lead to species-specific responses in the context of E2F4 recruitment in the developing brain. Overall, this research not only enhanced our understanding of the cellular and molecular mechanisms underlying radiation-induced microcephaly in humans but also highlighted the critical role of the DREAM complex in mediating p53-dependent gene repression. Moreover, it emphasizes the importance of employing in vitro human models in human disease contexts, allowing for the identification of species-specific mechanisms and paving the way for more accurate therapeutic approaches.

## 4. Materials and Methods Data and materials availability

All raw sequencing data can be accessed at EBI ArrayExpress (accession number: E-MTAB- 14150). Data from (23) can be accessed at NCBI Gene Expression Omnibus (accession number: GSE140464).

### Animals

Animal experiments were performed in accordance with the Ethical Committee Animal Studies of the Medanex Clinic (EC_MxCl_2014_036, EC_MxCl_2020_164), and in compliance with the Belgian laboratory animal legislation and the European Communities Council Directive of 22 September 2010 (2010/63/EU). All the mice were housed under standard laboratory conditions with 12 h light/dark cycle. Food and water were always available. Female and male mice were coupled during 2 hours in the morning, at the start of the light phase. The morning of coupling was considered E0.

### Cell culture

#### Human ESCs

The H7 (WA07) hESC line was obtained from WiCell (WiCell, #wa07, RRID:CVCL_9772). Cells were cultured on 35 mm dishes (Thermo Fisher, #150460) coated with Matrigel (Corning, #354277), and maintained in mTeSR Plus medium (Stem Cell Technologies, #100-0276) at 37 °C in a humidified atmosphere with 5% CO2. The cell culture medium was refreshed daily. Cells were free from mycoplasma infection. hESCs were used in accordance with the current ethical regulations.

#### Human ESC-derived NPCs

H7-derived hNPCs were generated following the technical manual from Stem Cell Technologies, with slightly modifications. Briefly, once 80% confluence was reached, hESCs were detached and disassociated into single cells using Accutase (Stem Cell Technologies, #07920). Next, 45.000 cells/well were transferred to ∼30 wells of a v-bottom 96-well plate (ThermoFisher, #277143) treated with Anti-Adherence Rising Solution (Stem Cell Technologies, #07010) that allows proper spheroid formation. Spheroids were maintained in SMADi Neural Induction Medium (Stem Cell Technologies, #08581) supplemented with 10 µM ROCK inhibitor Y-27632 (ROCKi) (Stem Cell Technologies, #72304) until Day 4. Partial medium change was performed daily. On Day 5, neurospheres were transferred to 35 mm dish coated with 15% Poly-L-Ornithine (Sigma-Aldrich, #P4957) and 10 µg/ml Laminin (Sigma-Aldrich, #L2020). Whole medium change was performed daily. On Day 8, successful neural induction was confirmed by the increased number of neural rosettes in culture. On Day 12, rosettes were selected by a 1.5-hour incubation with Neural Rosette Selection Reagent (Stem Cell Technologies, #05832), collected in a 15 ml tube, centrifuged at 300 g for 5 min, gently resuspended in SMADi Neural Induction Medium/ROCKi, and then NPCs at passage 0 were reseeded in one well of a 6-well plate coated with Poly-L-Ornithine and Laminin. After 2 days, SMADi Neural Induction Medium was replaced with Neural Progenitor Medium (Stem Cell Technologies, #05833). Cells were maintained at 37 °C in a humidified atmosphere with 5% CO2. The cell culture medium was refreshed daily. Cultures were free from mycoplasma infection.

#### Primary mouse NPCs

For RNA-seq primary mNPCs were derived from prefrontal cortex or medial ganglionic eminences (MGE) of embryonic day (E)13 fetuses. For qRT-PCR primary mNPCs were derived from prefrontal cortex of E15 wild-type or conditional p53 knockout (*Emx1*-Cre; *Trp53*^fl/fl^) (23) fetuses. Brain regions were separated and gently dissociated in Accutase (Stem Cell Technologies, #07920). NPCs were cultured as monolayers on Poly-D-Lysine coated 6-well plates (Corning, #356413), and maintained in Dulbecco’s modified eagle medium (DMEM)/F-12 (Gibco, #11330032) supplemented with 1% B-27 (Gibco, #17504044), 0.5% N-2 (Gibco, #17502048), 10 ng/ml of recombinant mouse EGF (PeproTech, # 315-09-1MG), and 20 ng/ml recombinant human FGF-basic (PeproTech, #100-18B). For MGE-derived mNPCs, EGF was omitted from the culture medium. Cells were incubated at 37 °C in a humidified atmosphere with 5% CO2. The cell culture medium was refreshed daily. Cells were free from mycoplasma infection.

#### Mouse embryonic fibroblasts

The STO MEF cells (ATCC, #CRL-1503, RRID:CVCL_3420) were cultured in DMEM (ATCC 30-2002) supplemented with 10% fetal bovine serum (Gibco, A5670701). Cells were incubated at 37°C in a humidified atmosphere with 5% CO2 and sub-cultured every three to four days when cells reached 70-80% confluence. Cells were free from mycoplasma infection.

#### Silencing of TP53 in hNPCs

For transient *TP53* knockdown, hNPCs were transfected with si-RNA for *TP53* (Dharmacon, ON-TARGETplus SMARTpool siRNA #L-003329-00-0005). Control samples were transfected with a non-targeting si-RNA (Dharmacon, ON-TARGETplus Non-targeting Control Pool, #D-001810- 10-05). hNPCs were transfected for 48 h with 5 nM siRNA using Lipofectamine RNAiMAX Transfection Reagent (Thermo Fisher Scientific, #13778075).

#### HLM006474 (E2F inhibitor) treatment

HLM006474 (MedChemExpress, #HY-16667) was prepared in dimethyl sulfoxide. hNPCs, mNPCs (wild-type and p53 knockout), and MEFs were treated with 10 µM or 40 µM of HLM006474, while control cells received an equal concentration of dimethyl sulfoxide, never exceeding 1%. Both control and treated cells were exposed to IR 6 h following treatment and were lysed 24 h after irradiation for RNA extraction.

### Generation of forebrain organoids

Forebrain organoids were generated as previously described (45), with minor modifications. Briefly, once 80% confluence was reached, hESCs were detached and disassociated into single cells using Accutase. Next, 3.000.000 cells were transferred to a well of an AggreWell™ 800 plate (Stem Cell Technologies, #34811) treated with Anti-Adherence Rising Solution. Cells were maintained in mTeSR Plus medium supplemented with 10 µM ROCKi for 24 hours. On the next day, spheroids were resuspended, filtered using a 40 µm strainer (Corning, #352340), and transferred to low attachment T75 flasks (ThermoFisher, #174952) containing Neural Induction Medium (DMEM F/12 (Gibco, #11330032), 20% knockout serum replacement (Gibco, #10828028), 1% MEM non-essential amino acids (Gibco, #11140050), 0.5% GlutaMAX (Gibco, #35050061), 0.1 mM 2-mercaptoethanol (Sigma-Aldrich, #M3148), 1% penicillin/streptomycin (Sigma-Aldrich, #P4333), and 0.1% MycoZap (Lonza Group, #VZA-2031)) supplemented with 5 µM Dorsomorphin (Sigma-Aldrich, #P5499) and 10 µM SB-431542 (Tocris Bioscience, #1614). Medium change was performed daily until Day 5. On Day 6 of culture, Neural Induction Medium was replaced by Neural Differentiation Medium (Neurobasal-A Medium (Gibco, #10888022), 2% B-27 supplement without vitamin A (Gibco, #12587010), 0.5% GlutaMAX, 1% penicillin/streptomycin, and 0.1% MycoZap) supplemented with 20 ng/ml mouse recombinant EGF (PeproTech, # 315-09-1MG) and 20 ng/ml human recombinant FGF-basic (PeproTech, #100- 18B) until Day 24, which was then replaced by 20 ng/ml human/mouse recombinant NT-3 (Stem

Cell Technologies, #78074) and 20 ng/ml human recombinant BDNF (Stem Cell Technologies, #78005) until Day 43 of culture. Half medium change was performed every other day. From Day 43 on, growth factors were removed, and whole medium change was performed every 4 days. Organoids were kept in culture for a minimum of 14 days and a maximum of 70 days. Organoid cultures were maintained at 37 °C in a humidified atmosphere with 5% CO2, and were free from mycoplasma infection.

### BrdU labelling of forebrain organoids

The thymidine analogue BrdU (Sigma-Aldrich, #B5002) was used to label proliferative cells at the time of IR exposure. Briefly, forebrain organoids at developmental days D14 and D56 were treated with 100 µM BrdU for 2 h and then washed 3x with DMEM/F12. Next, samples were irradiated and subsequently fixed after 24 h and 14 days, following the procedures described in the *Ionizing Radiation (IR) Exposure* and *Immunostaining* sections of this manuscript, respectively.

### Ionizing radiation (IR) exposure

Samples were given a single dose of X-ray using an X-Strahl 320 kV machine (250 kV, 12 mA, 3.8 mm AI equivalent, 1.4 mm Cu-filtered X-rays) in accordance with ISO 4037. Irradiation was performed vertically at a distance of 100 cm, with samples being placed within the range of the X-ray beam to ensure homogeneity. Human forebrain organoids received doses of 0.5 Gy or 2 Gy, while hNPCs and mNPCs received 1 Gy. Control samples were taken to the radiation facility but were not placed within the radiation field (sham-irradiation or 0 Gy). Samples were processed after 2 h, 6 h, 24 h or 14 days upon IR exposure. For each irradiation-related experiment, at least three biological replicates of forebrain organoids, and two to four biological replicates of mNPCs, and two to four technical replicates of hNPCs were used, as indicated in figure legends.

### Human forebrain organoid size determination

During development, forebrain organoid size was measured every 2 days. Following IR exposure, measurements were taken every 2 days for 14 days. Samples were imaged under an inverted Leica DMi1 microscope (Leica Microsystems, Germany). Due to size limitations, organoids were imaged under a stereo Leica MZ12 microscope (Leica Microsystems, Germany) after Day 45 of culture. The surface diameter was measured using ImageJ, with 24 organoids per condition being analyzed.

### Measurement of neural rosette perimeter and thickness

Neural rosette thickness was measured at four distinct locations (T1, T2, T3, and T4), and the average thickness was then determined using the formula: Thickness (T) = (T1 + T2 + T3 + T4) / 4 (Figure S3C). The perimeter of the neural rosette was measured using the elliptical tool in the ImageJ software, which allows for precise tracing of the rosette’s boundary.

### Immunostaining

Organoids were washed once with warm DMEM/F12, transferred to 2 ml tubes (1 per tube), and fixed in 4% PFA for 30 min at room temperature (RT). Samples were washed 2x with phosphate buffered saline (PBS). Fixation was performed 2 h, 6 h, 24 h and 14 d following irradiation. Next, organoids were cryopreserved in 30% sucrose dissolved in PBS overnight, and embedded in Neg-50 Frozen Section Medium (Epredia, #6502). Organoids were sectioned into 10-μm thick cryosections (ThermoFisher, CryoStar NX50) and mounted on SuperFrost Plus Adhesion Slides (Epredia, #J1800AMNZ). Before staining, cryosections were incubated in acetone for 2 min, air-dried for 30 min, washed in dH2O for 5 min, and boiled 2x for 4:30 min at 700W and 1x for 5:00 min at 600W in citrate buffer, pH 6.1 (Dako, #S1699) for antigen retrieval. Sections were cooled down at RT for 20 min. Next, samples were washed 1x for 5 min with PBS, permeabilized 3x for 5 min with 0.25% Triton X-100 in tris-buffered saline (TBS/Triton). The blocking of nonspecific bindings was performed either with 5% Bovine Serum Albumin or pre-immune goat serum (1:5) in Tris-NaCl buffer for 1 hr at RT. Next, sections were incubated with primary antibodies diluted in blocking solution overnight at 4 °C. The following antibodies were used: anti-SOX2 (rabbit, 1:300 (Abcam, #ab97959)), anti-Nestin (mouse, 1:300 (Invitrogen, #MA1-110)), anti-PAX6 (rabbit, 1:200 (BioLegend, #901301)), anti-DCX, (mouse, 1:300 (Santa Cruz, #sc271390)), anti- TUJ1 (rabbit, 1:300 (Sigma Aldrich, #T2200)), anti-ɣH2AX (mouse, 1:200 (Millipore, #JBW301)), anti-53BP1 (rabbit, 1:200 (Novus Biologicals, #NB100-304)), anti-p-p53 (rabbit, 1:200 (Abcam, #ab1431)), anti-CC3 (rabbit, 1:500 (Cell Signaling, #9661)), anti-Ki67 (rabbit, 1:500 (Abcam, #ab15580)), anti-pVimentin (mouse, 1:300 (Abcam, #ab22651)), anti-PH3 (rabbit, 1:200 (Cell Signaling, #3377), anti-BrdU (rat, 1:300 (Abcam, #ab6326)). The following day, samples were washed 3x for 5 min with TBS/Triton and incubated 2 h with the appropriate Alexa Fluor-488, -568, -555, or Cy5 (Invitrogen) secondary antibody diluted 1:200 in blocking solution at RT. Sections were then washed 3x for 5 min with TBS/Triton, counterstained with DAPI for 15 min, washed 1x for 5 min with TBS/Triton and 1x with dH2O. Finally, samples were mounted with

ProLong Diamond Antifade Mountant (Invitrogen, #P36962), air-dried overnight at RT, and stored at 4 °C until imagining.

For Ki67 and CC3 stainings, signal amplification was performed. Sections were incubated with the appropriate anti-HRP secondary antibody (Dako) diluted 1:200 in blocking solution for 2 h at RT. Next, samples were washed 3x for 5 min with TBS/Triton and incubated with TSA Plus Cyanine 3 (PerkinElmer, #NEL744001KT) for 8 min at RT.

Terminal deoxynucleotidyl transferase (TdT) dUTP Nick-End Labeling (TUNEL) assay was performed using the Click-It Plus TUNEL for In Situ Apoptosis Detection, Alexa Fluor™ 488 dye (ThermoFisher, #C10617), as described in the supplier’s data sheet.

### Immunofluorescence microscopy

Samples were imaged using 20x, 40x or 60x objectives either on a Nikon Eclipse Ti-E inverted microscope (Nikon, Japan) or a Leica SP8 confocal microscope (Leica Microsystems, Germany). On the confocal, images were always acquired as Z-stacks with a 0.5 µm splicing interval and with an image resolution of 1024 x 1024 pixels. Pictures were analyzed using ImageJ.

### RNA library preparation for RNA sequencing (RNA-Seq)

Human forebrain organoids were irradiated and samples were lysed at 6 h, 24 h and 14 d following IR exposure. Organoid lysis was performed in RLT Plus lysis buffer (Qiagen, #74136) and samples were stored at -80°C until RNA extraction using the RNeasy Mini Kit (Qiagen, #74136). RNA quality was determined using the 5400 Agilent fragment analyzer system (Agilent Technologies, United States). Most of the samples had RNA integrity numbers >9.0. cDNA library construction and sequencing were performed by NovoGene (Cambridge, United Kingdom). Messenger RNA was purified from total RNA using poly-T oligo-attached magnetic beads. After fragmentation, the first strand cDNA was synthesized using random hexamer primers, followed by the second strand cDNA synthesis using either dUTP for directional library or dTTP for non-directional library. For the non-directional library, it was ready after end repair, A-tailing, adapter ligation, size selection, amplification, and purification. The library was checked with Qubit and real-time PCR for quantification and bioanalyzer for size distribution detection. Quantified libraries were pooled and sequenced (PE150) on Illumina platforms.

RNA-seq analysis was performed with nf-core/rnaseq version 3.12.0 (103) using the default values except for: i) Ensembl GRCh38 release 109 was used as the reference genome, ii) salmon quant was run with the “--gcBias” parameter.

Differential expression analysis was performed with nf-core/differentialabundance version 1.4.0 (103) with the default values. For analyses of effects of developmental timing, sham-irradiated (0 Gy) organoids of D14 + 6h, D14 + 14 days, D56 + 6 h, and D56 + 14 days were considered as D14, D28, D56 and D70, respectively. For analyses of radiation effects, irradiated organoids (0.5 Gy or 2 Gy) were always compared to their respective sham-irradiated control at the same time point. Genes with a false discovery rate (FDR) <0.05 were considered differentially expressed.

### Deconvolution analysis

Deconvolution of bulk RNA-seq data was performed with BisqueRNA (104) version 1.0.5 in marker-based mode. The markers for the different cell types/states used were: Proliferation: MKI67, CENPF, TOP2A, NUSAP1, UBE2C; Astrocytes: GFAP, S100B, AQP4, GJA1; Oligodendrocytes: OLIG1, OLIG2, PDGFRA; Choroid plexus: TTR, AQP1, RSPO2, PLS3; Intermediate progenitors: EOMES, NHLH1, PPP1R17; Glutamatergic neurons: SLC17A6, SLC17A7, GRIN2B, NRN1; GABAergic: GAD1, GAD2, SLC32A1, NKX2-1, DLX1, DLX6-AS1; Dopaminergic neurons: TH, DDC, EN1.

### Statistical overlap analysis between categorical and ranked lists of DEGs

For comparisons between two categorical gene lists, we used hypergeometric tests. Colors of each square indicate -Log10(p-value) of the overlap, with a maximum set at 300.

Rank–rank hypergeometric overlap (RRHO) analysis was performed with the R package RedRibbon version 1.1.1 (105) to detect the overlap between genes differentially expressed in the same or opposite directions. Bottom left and top right quadrants in each map display overlap of shared down- or upregulated gene, respectively. The top left and bottom right quadrants display discordant overlaps.

### Gene Ontology and Transcription Factor binding site enrichment analysis

For analysis of overrepresented biological processes and TF binding sites we used the “GO Biological Process 2023” and “ENCODE and ChEA Consensus TFs from ChIP-X” datasets within Enrichr (https://maayanlab.cloud/Enrichr/) (106) and Metascape for simultaneous enrichment analysis (GO/KEGG terms, canonical pathways, and MSigDB Hallmark gene sets) of multiple gene lists (107). Networks from Metascape were adapted using Cytoscape v3.10.1 (108).

### Gene Regulatory Network analysis: module generation and eigengene expression

Length-scaled counts were loaded into R (v.4.1.2). They were preprocessed using the R package edgeR (109) (R v.4.1.2) by filtering genes on a counts per million (cpm) value greater than one in at least two samples, normalizing by library size (110), and logarithmic transformation with a prior count of one. Next, highly variable, protein-coding genes were selected, after which additional TFs and microcephaly genes were added (respectively, based on Lovering *et al.* (111) and DisGeNET (112): C0431350, C3711387 and C1855081). The scaled count data of these 9434 genes was finally used as input for module network inference by Lemon-Tree (in Java as a command-line program (v3.1.1), https://github.com/erbon7/lemon-tree) (113). In a first step, ensemble clustering (100 times) inferred 200 coexpression modules. In a next step, regulators were assigned to the modules based on TF activity for 678 TFs as predicted using the R package decoupleR (114) with the most recent version (Dec. 2023) of CollecTRI (115) as a prior network. The cut-off of the regulator scores were taken as was set two times the highest random score (6.5). This resulted in 323 unique TFs assigned to the modules. Motif enrichment analysis was performed with RcisTarget in R (116) version 10 in genomic regions 10 kb up- and downstream of the TSS and the files were downloaded from https://resources.aertslab.org/cistarget/.

Average expression plots of the modules were made with ggplot2 in R. The expression was averaged for each sample over all genes in one module. The full line represents the average expression while the dotted lines represent the Q1 and Q3 values.

### Gene regulation by p53 and E2F4/DREAM

Human and mouse MCPH genes were evaluated for their regulation by p53 and E2F4 (for mouse genes) or DREAM (for human genes) based on their p53 Expression Scores and E2F4 or DREAM Binding Scores from (75). E2F4 ChIP-seq read counts at the promoters of *CIT*, *KNL1*, *ASPM* and *WDR62* were derived from www.ChIP-Atlas.org (74).

### Quantitative reverse transcriptase PCR (qRT-PCR)

Human forebrain organoids, hNPCs, and mNPCs were irradiated and samples were lysed following IR exposure (Organoids: 6 h, 24 h and 14 d after IR exposure; mNPCs/hNPCs: 6 h, 24 h after IR exposure). Sample lysis was performed in RLT Plus lysis buffer and samples were stored at -80°C until RNA extraction. RNA was extracted following the Qiagen RNEasy Mini kit protocol, and eluted in 30 μl of RNase-free water. Next, cDNA was synthetized using the GoScript Reverse Transcriptase kit (Promega, #A2801). qRT-PCR was then performed using a qTOWER³ G machine (Analytik Jena, Germany) and the QuantiNova SYBR Green (Qiagen). Relative gene expression was calculated via the Pfaffl method (117) using *RPL13A* or *Polr2a* as reference genes. The list of primers can be found in Table S3. For all qRT-PCR experiments the specificity of the primers was validated using a melting curve.

### Statistical analysis

Statistical analysis was performed using GraphPad Prism 8 or 10. Comparisons between sham and irradiated samples were conducted. One-Way ANOVA test followed by Tukey’s test for multiple comparisons was performed when comparing between three conditions. A two-tailed Student’s t-test was performed when comparing between two conditions. A p-value of <0.05 was considered statistically significant. Further statistical details can be found in figure legends.

## Author contributions

Conceptualization: RQ, NR, JHR; Methodology: JHR, RQ, EE, LB, VV, HP; Investigation: JHR, JB, AJ, LB, ACMM; Data curation: EE; Formal Analysis: JHR, RQ, EE, HP, VV; Visualization: JHR, RQ, EE, HP; Resources: WHDV; Funding acquisition: RQ, SB, WVDH; Project administration: RQ, SB; Supervision: RQ, NR, SB; Writing – original draft: JHR, RQ; Writing – review & editing: NR, WHDV, VV, LB, HP, ACMM.

## Funding

The author(s) declare financial support was received for the research, authorship, and/or publication of this article. This work was supported by the Research Fund Flanders (G0A3116N to RQ). JR and LB are the recipient of a SCK CEN PhD scholarship. JR is the recipient of a Special Research Fund (BOF/UGent) for a PhD trajectory that was delayed due to COVID-19 (BOF23/CDV/144). JR was granted a MELODI mobility grant and a PIANOFORTE travel grant for early career researchers. The Leica SP8 confocal system, was funded by the Hercules Foundation and the Flemish Government (GOH4216N to WHDV).

### Acknowledgments

The authors would like to acknowledge ACAM, the microscopy core facility of the University of Antwerp, and thank Isabel Pintelon for her assistance during the microscopy sessions. We thank Lara Barazzuol and Daniëlle Voshart (University Medical Center Groningen) for valuable feedback for organoid generation, and Laura Pellegrini (King’s College London) for valuable discussions.

## Conflict of interest

The authors declare that the research was conducted in the absence of any commercial or financial relationships that could be construed as a potential conflict of interest.

---

**Figure S1.**
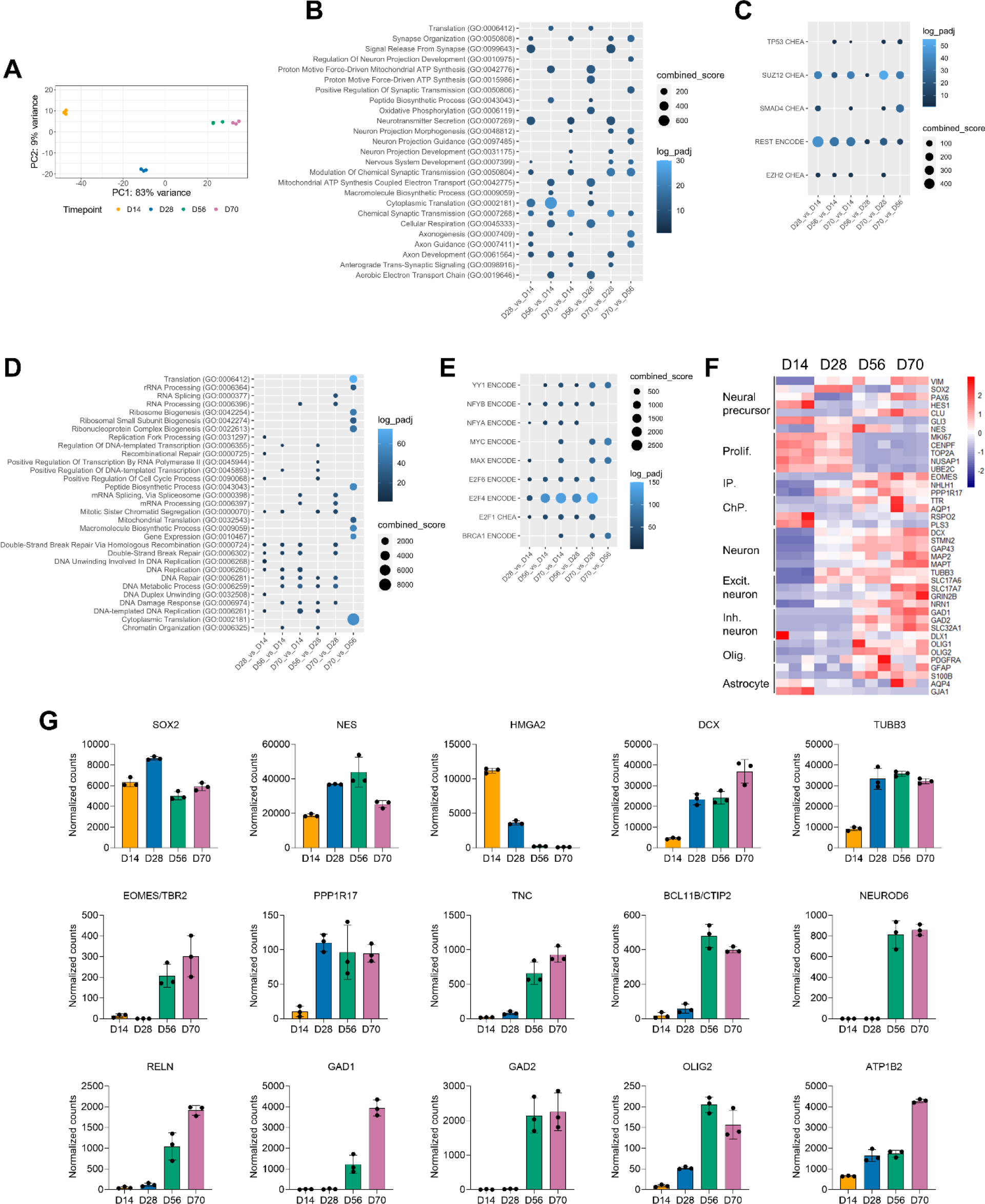
Gene ontology and transcription factor enrichment of developmentally up- and downregulated genes. (A) Principal component analysis of D14, D28, D56 and D70 organoids. (B-E). Bubble plots showing enriched GO Biological Processes (B, D) and predicted transcriptional regulators (C, E) of differentially (FDR <0.05) upregulated genes (B, C) and downregulated genes (D, E) for each indicated pairwise comparison. F. Heatmap showing vst- normalized and row-scaled expression levels of marker genes. Normalized (by variance-stabilized transformation, vst) RNA-seq count data of different brain cell type markers across different stages of organoid development (D14, D28, D56, D70).

**Figure S2.**
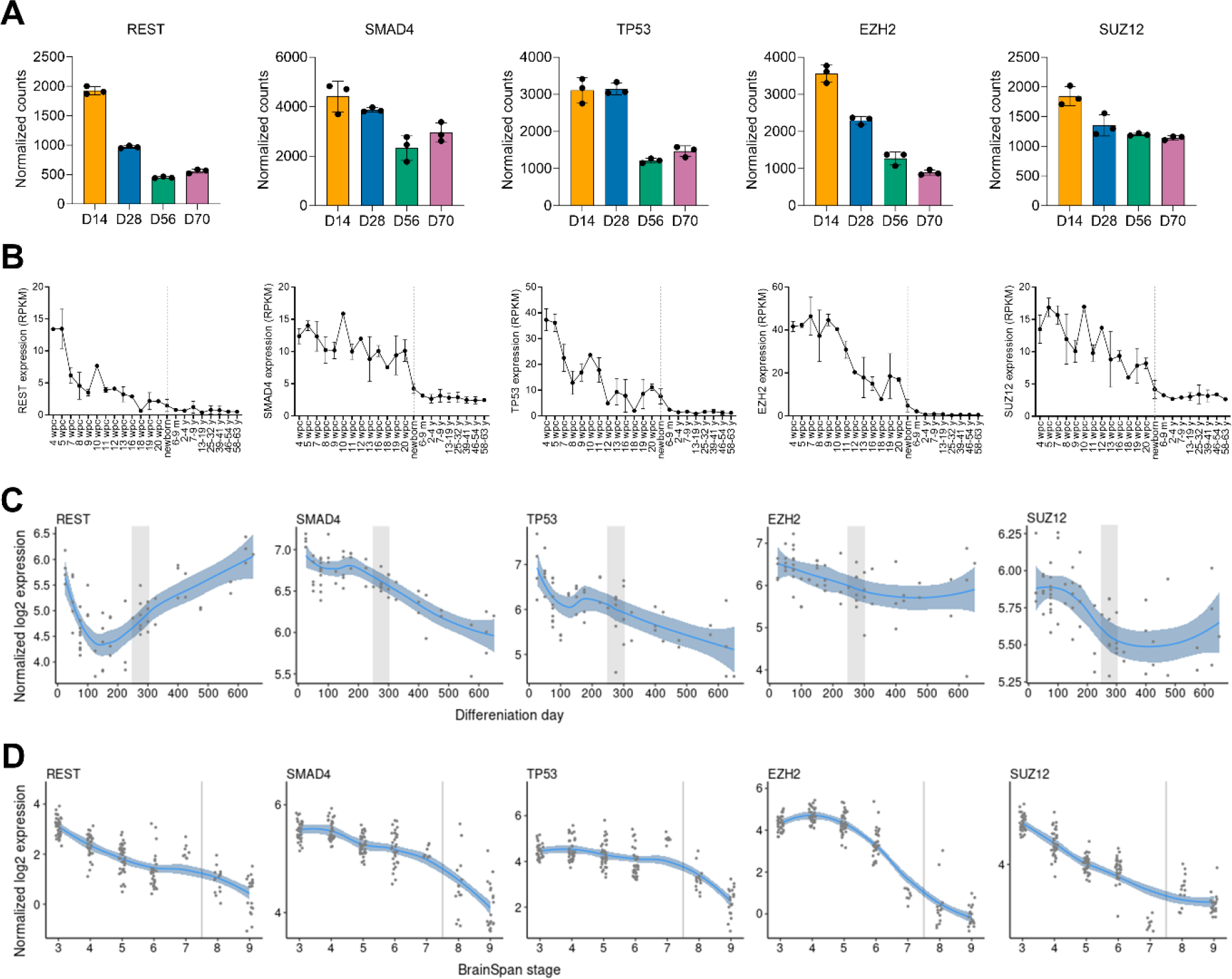
Developmental gene expression profiles of transcription factors predicted to regulate developmentally enriched genes. (A) Vst-normalized gene expression profiles in organoids across different stages of organoid development (D14, D28, D56, D70) from this study. (B) Gene expression in human brain tissue at different pre- and post-natal stages. The vertical line indicates birth. Data downloaded from https://apps.kaessmannlab.org/evodevoapp/ (52). (C, D) Gene expression in human forebrain organoids (C) and Brainspan (D) Data plotted from http://solo.bmap.ucla.edu/shiny/GECO/ (53).

**Figure S3.**
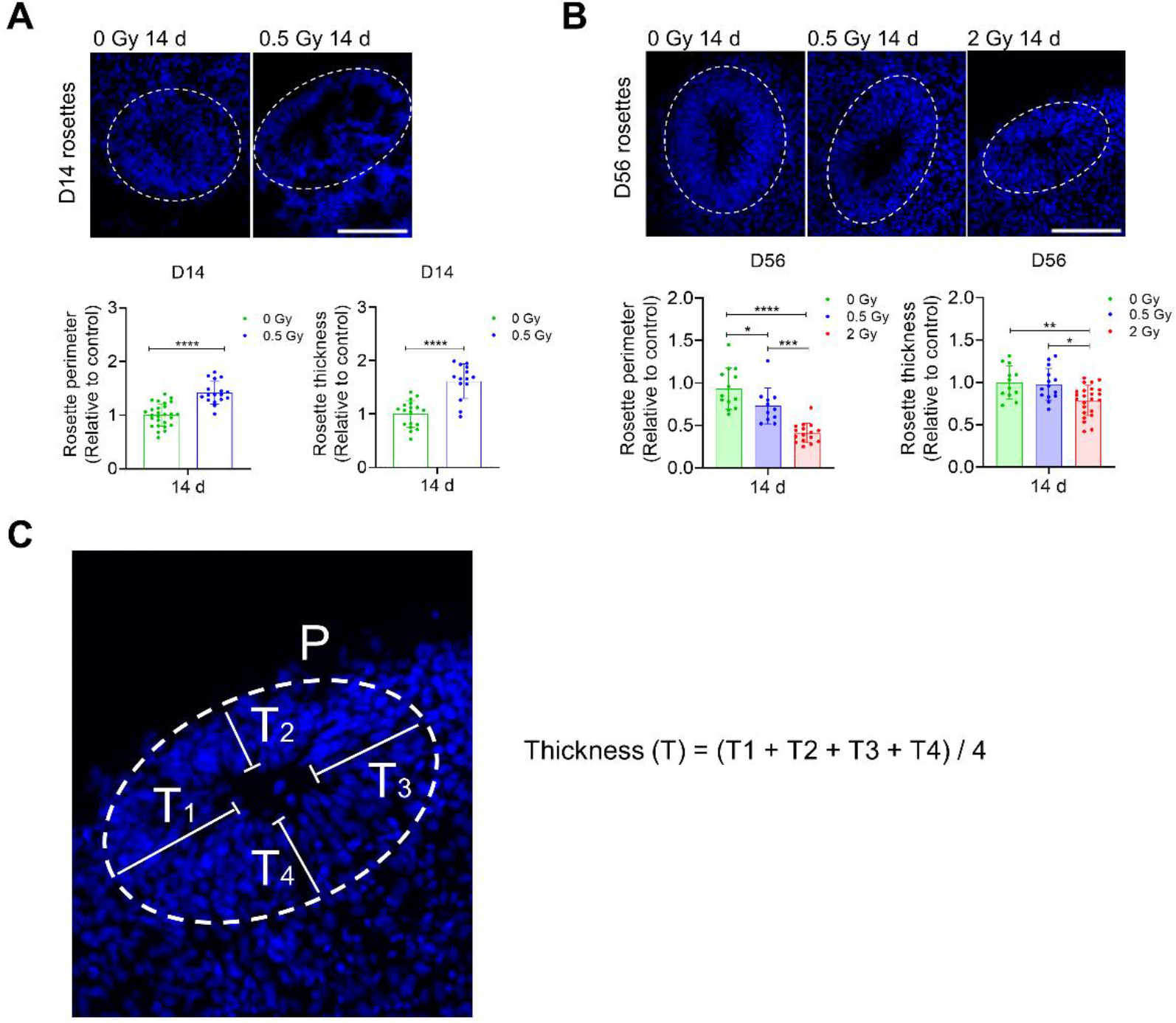
Radiation-induced changes in neural rosette perimeter and thickness. (A, B) Neural rosette perimeter and thickness were measured. Four organoids were analyzed per condition. Image analysis was performed using the ImageJ software. One-way ANOVA test followed by Tukey’s test for multiple comparisons was used when comparing 3 conditions, *p<0.03 **p< 0.005; ***p< 0.0003; ****p< 0.0001. Student’s t-test was used when comparing 2 conditions, ****p< 0.0001. Scale bar = 100 µm. (C) Neural rosette thickness was measured at opposite sides of the primary and secondary axes (T1, T2, T3, and T4), and the average thickness was then determined using the formula: Thickness (T) = (T1 + T2 + T3 + T4) / 4.

**Figure S4.**
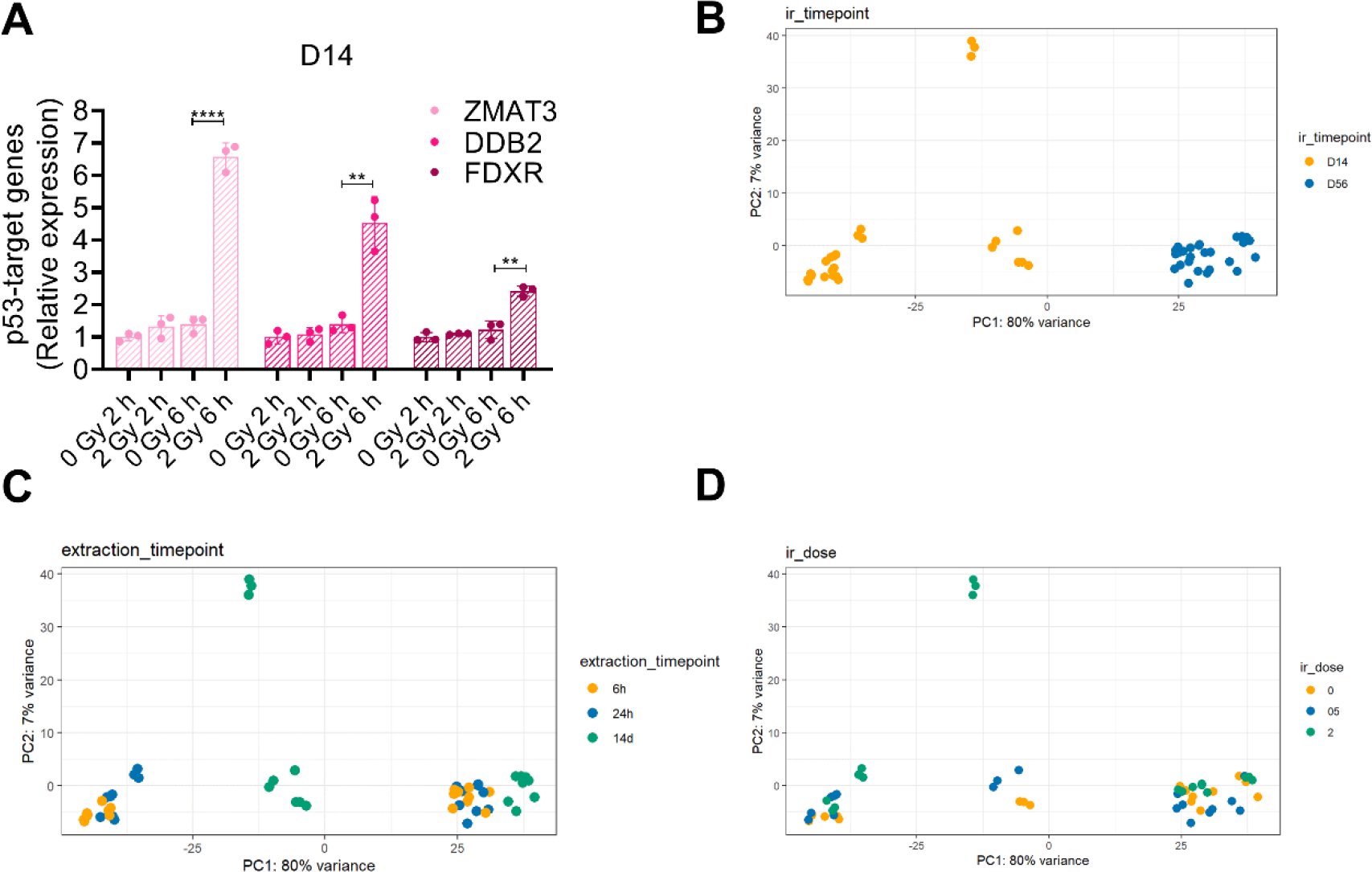
qPCR of selected p53 targets and principal component analysis plots of RNA-seq data. (A) P53 target genes were not yet induced at 2 h following exposure. n = 3. Student’s t-test was used,**** p<0.0001, **p< 0.003. (B-D) PCA plots of 54 experimental samples indicating effect of irradiation time point (B), RNA extraction time point (C), and irradiation dose (D).

**Figure S5.**
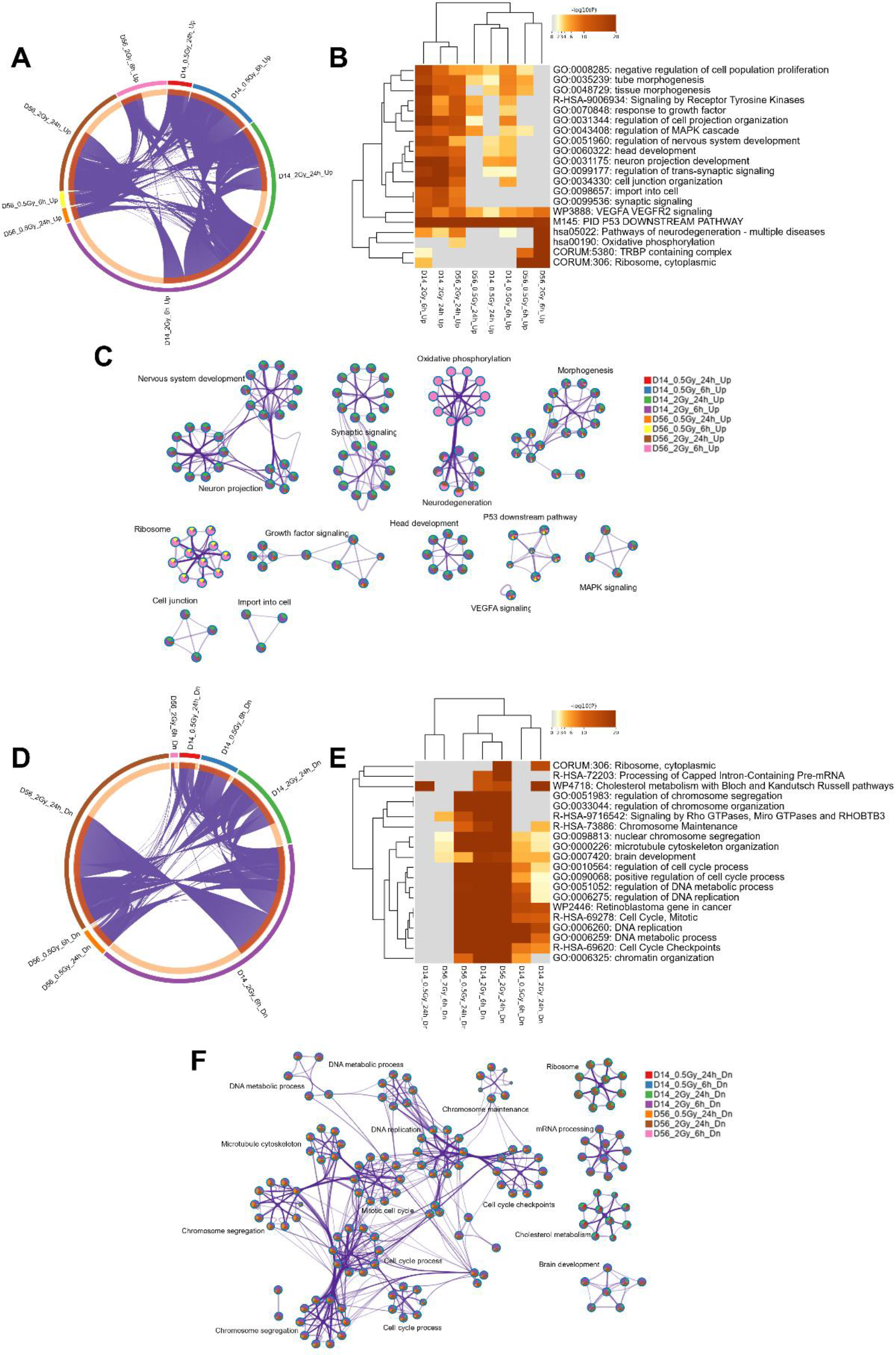
Functional enrichment analysis of early response (6 h, 24 h) to radiation using Metascape. (A) Circos plot indicating overlapping upregulated genes between indicated comparisons. Genes shared among gene lists are linked via purple lines. (B) Heatmap showing hierarchical clustering of top enriched functional categories for upregulated genes. Colors indicate level of significance. Grey means no significant enrichment. (C) Enrichment network visualization, where node colors indicate association with each input gene list. (D-F) The same as A-C, for downregulated genes.

**Figure S6.**
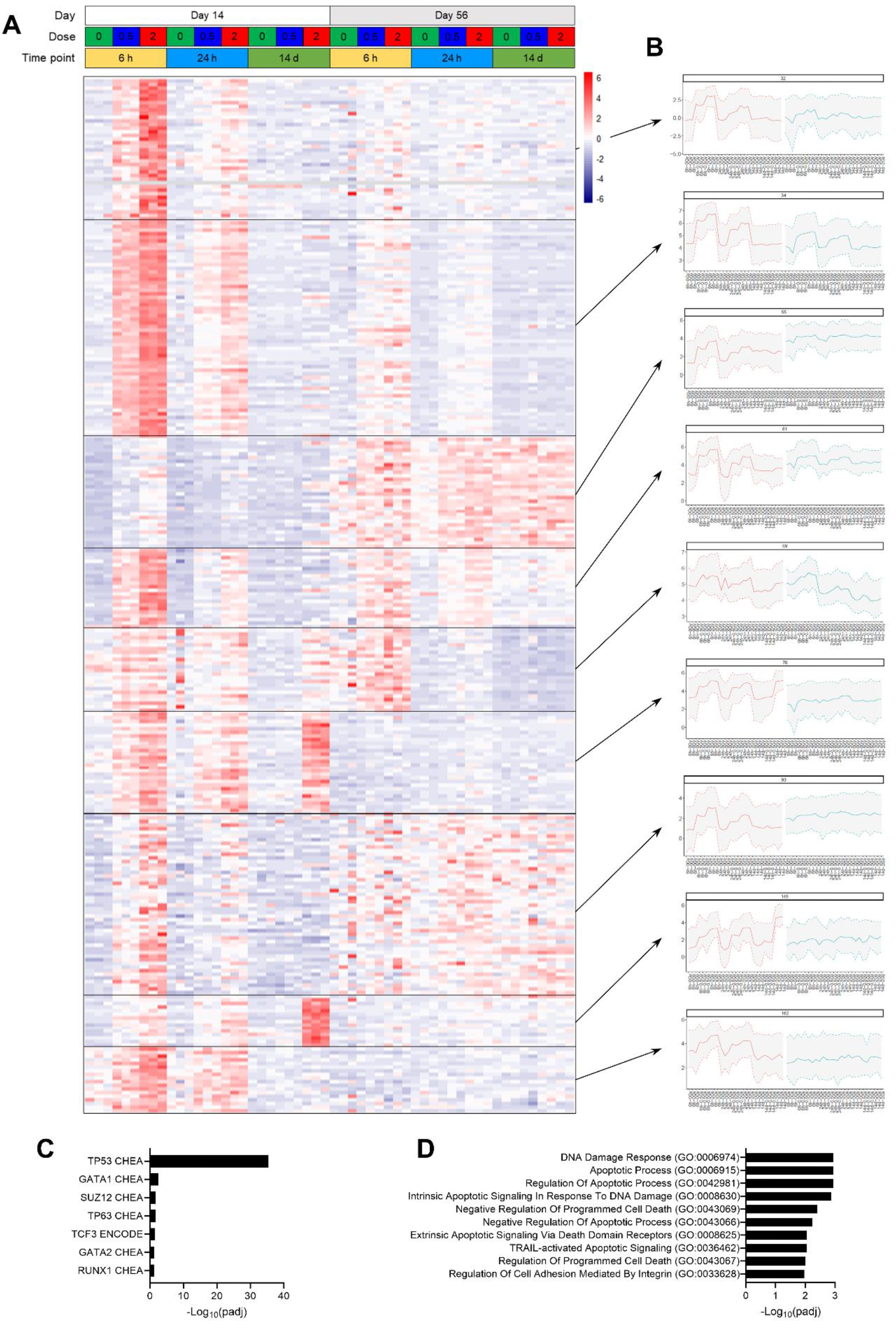
Modules of early (6 h, 24 h) dose-dependently upregulated genes are related to the classical p53-dependent DDR. (A) Heatmap showing normalized (by variance stabilizing transformation, vst) and row-scaled expression levels of genes belonging to nine modules of upregulated genes. (B) Eigengene expression profiles for each module, with interquartile ranges. Samples were ranked as in the heatmap in (A). (C-D) TF and GO-BP enrichment according to Enrichr.

**Figure S7.**
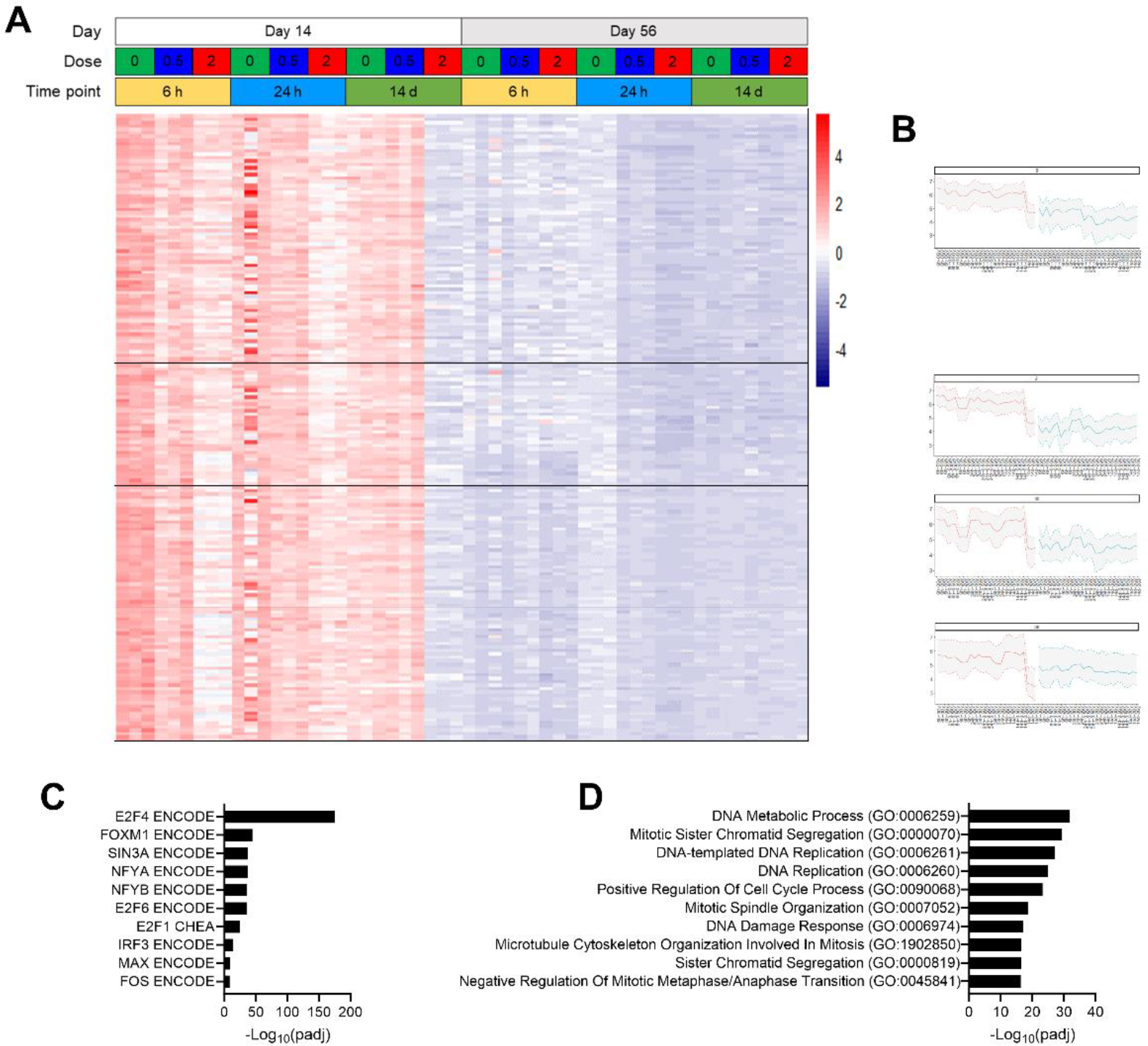
Modules of early (6 h, 24 h) dose-dependently downregulated genes are related to E2F4-dependent processes. (A) Heatmap of genes belonging to 4 modules of downregulated genes. (B) Eigengene expression profiles for each module, with interquartile ranges. Samples were ranked is in the heatmap in (A). (C-D) TF and GO-BP enrichment according to Enrichr.

**Figure S8.**
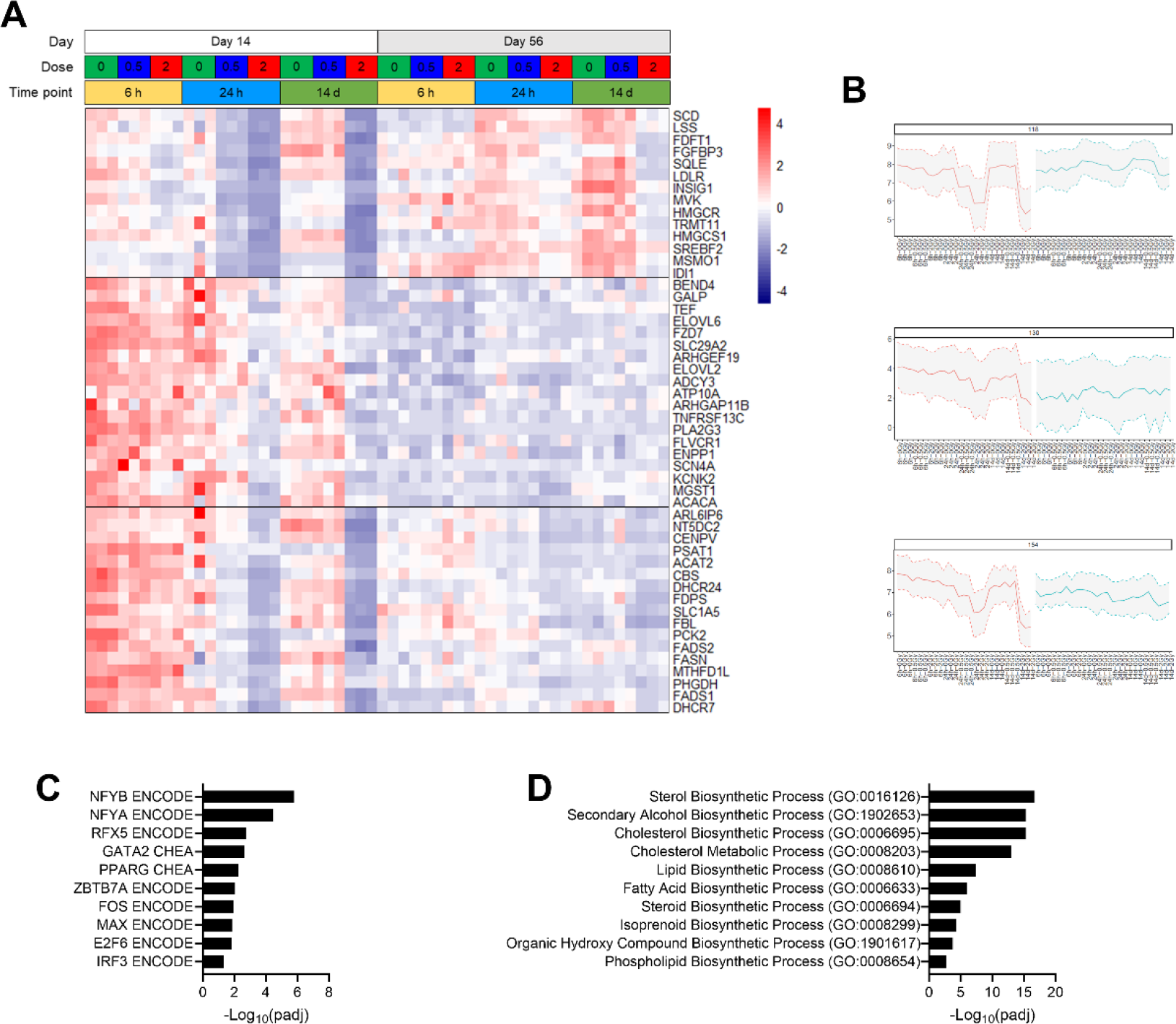
Modules of dose-dependently downregulated genes at 24 h after IR are related to NFY-dependent metabolic processes. (A) Heatmap showing vst-normalized and row-scaled expression levels of genes belonging to 3 modules of downregulated genes. (B) Eigengene expression profiles for each module. (C-D) TF and GO-BP enrichment according to Enrichr.

**Figure S9.**
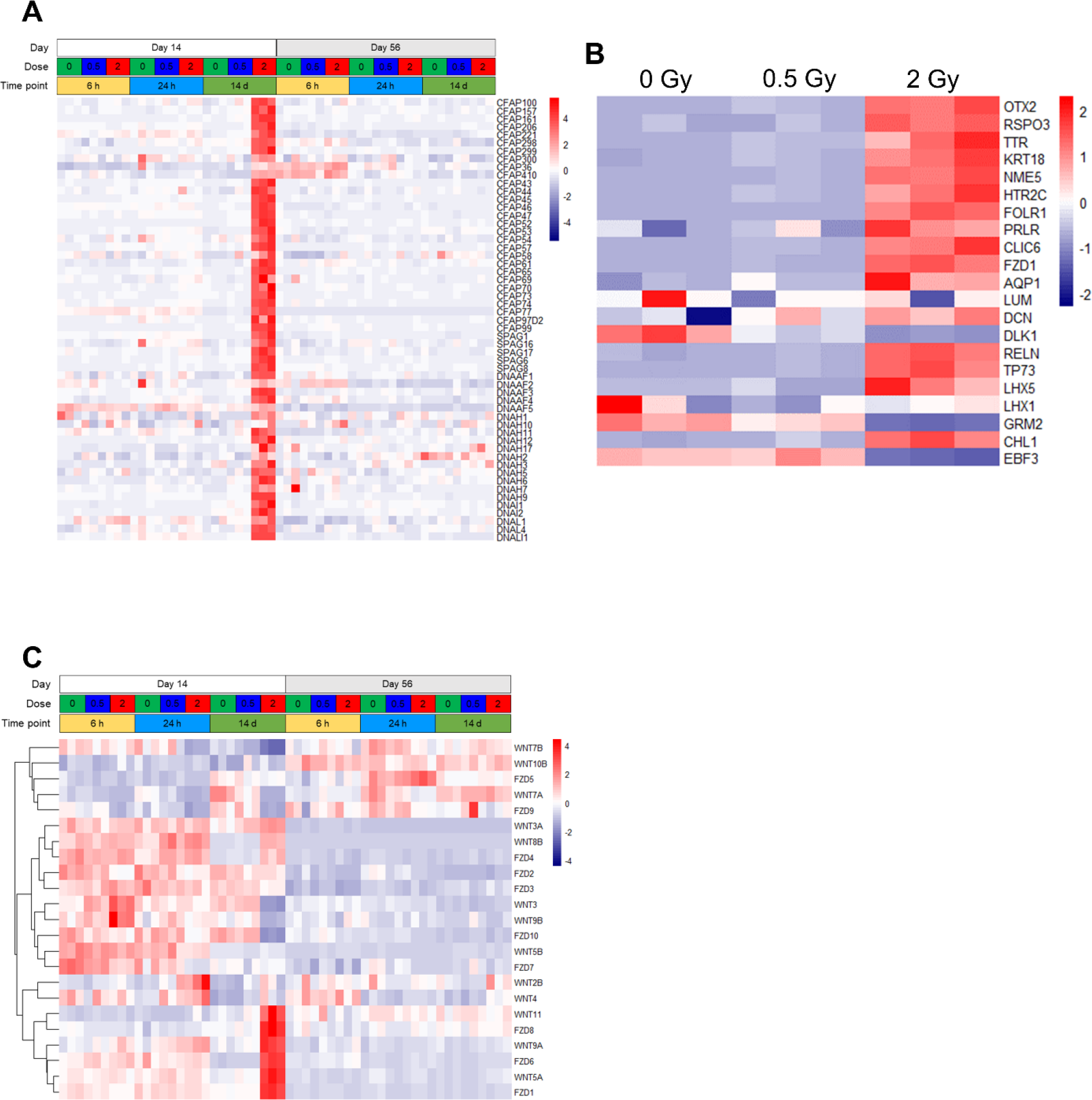
Markers of choroid plexus and Cajal-Retzius cells are severely dysregulated 14 days following 2-Gy exposure of D14 organoids. (A) Heatmap showing vst-normalized and row-scaled expression levels of genes related to cilia motility. (B) Heatmap showing vst-normalized and row- scaled expression levels of ChP and CR marker genes in D14 organoids at 14 days following irradiation. (C) Heatmap showing vst-normalized and row-scaled expression levels of WNT and FZD genes. Several of these genes are either highly up-or downregulated in 2-Gy irradiated D14 organoids.

**Figure S10.**
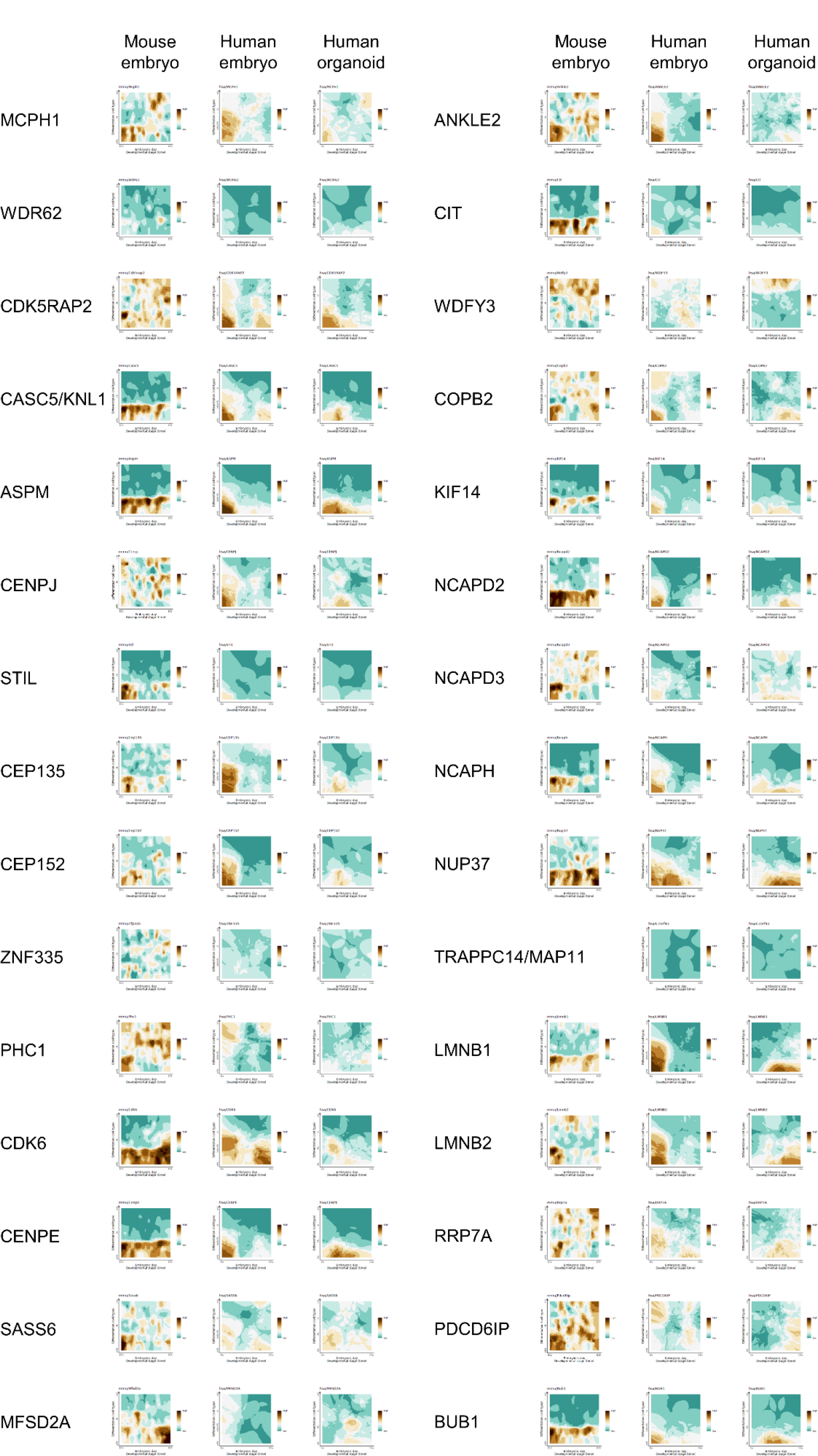
Spatiotemporal expression of MCPH genes in mouse and human embryonic brain and human brain organoids. Plots indicate developmental gene dynamics in specific cell types based on single-cell sequencing data from mouse embryonic brain (left), human embryonic brain (middle) and human brain organoids (right). Data were plotted from http://genebrowser.unige.ch/humous/ (62). Note that expression data for mouse Trappc14/Map11/BC037034 are not available. aRG, apical radial glia; bRG, basal radial glia; iN, immature neurons; IPC, intermediate progenitor cells; mN mature neurons.

**Figure S11.**
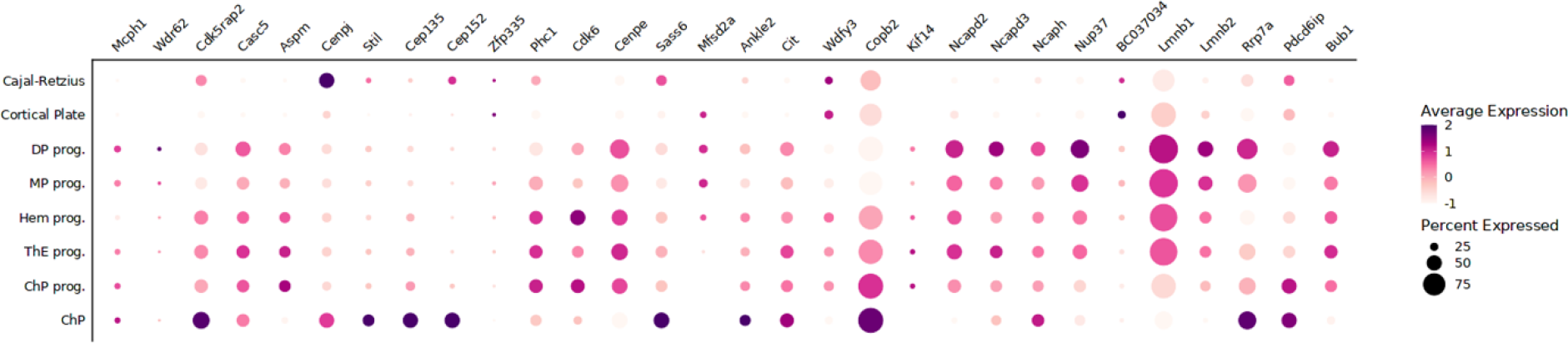
Bubble plot showing MCPH gene expression levels between different cell types around the early cortical hem in mice. Data were plotted from https://apps.institutimagine.org/mouse_hem/ (66).

**Figure S12.**
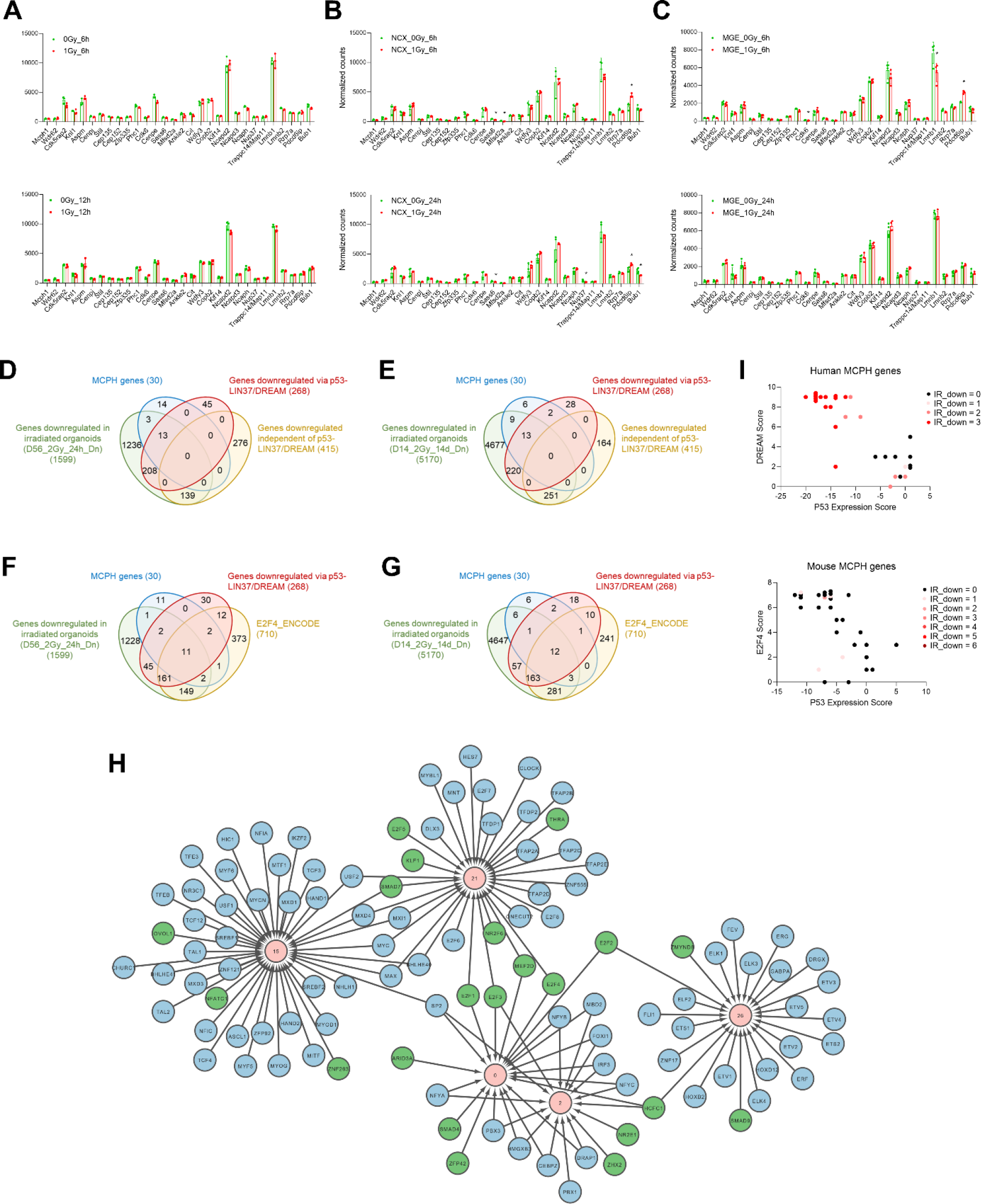
Irradiation does not generally affect MCPH gene expression in mouse models of radiation-induced microcephaly. (A-C) Normalized RNA-seq counts in 0-Gy (green bars) and 1- Gy (red bars) irradiated mouse embryonic brains (from (23)) (A), primary mouse cortical NPCs (B), and primary mouse MGE-derived NPCs (C). *denotes significant differential expression (FDR <0.05) in comparison to 0 Gy. (D-G) Overlaps between repressed genes in D56 organoids after 24 h (D, F), D14 organoids after 14 days (E, G), MCPH genes, targets of p53-LIN37/DREAM and p53-repressed LIN37/DREAM independent genes (D, E), or E2F4 targets (F, G). (H) Network of gene modules enriched in MCPH genes with their regulatory TFs. The modules are depicted in pink, while the TFs are in green or blue. TFs inferred by LemonTree are in green, while TFs found only with motif enrichment are in blue. Only unique TFs for each module are depicted, hence they are in green if they were found by both LemonTree and the motif enrichment. The top ten (according to NES) motif enrichment TFs are depicted. If multiple motifs had the same NES score as the tenth ranked motif, those additional motifs were included as well, resulting in potentially more than ten TFs. The microcephaly modules are co-regulated by some TFs, such as E2F factors, HCFC1, SMAD7, MEF2D, and MYC. Modules 0 and 2 share most TFs. (I) p53 Expression Score plotted against DREAM Binding Score (left) or E2F4 Binding Score (right). Dots are colored according to the number of conditions in which they were downregulated in human organoids (left) or mouse embryonic brain or NPCs (right).

**Figure S13.**
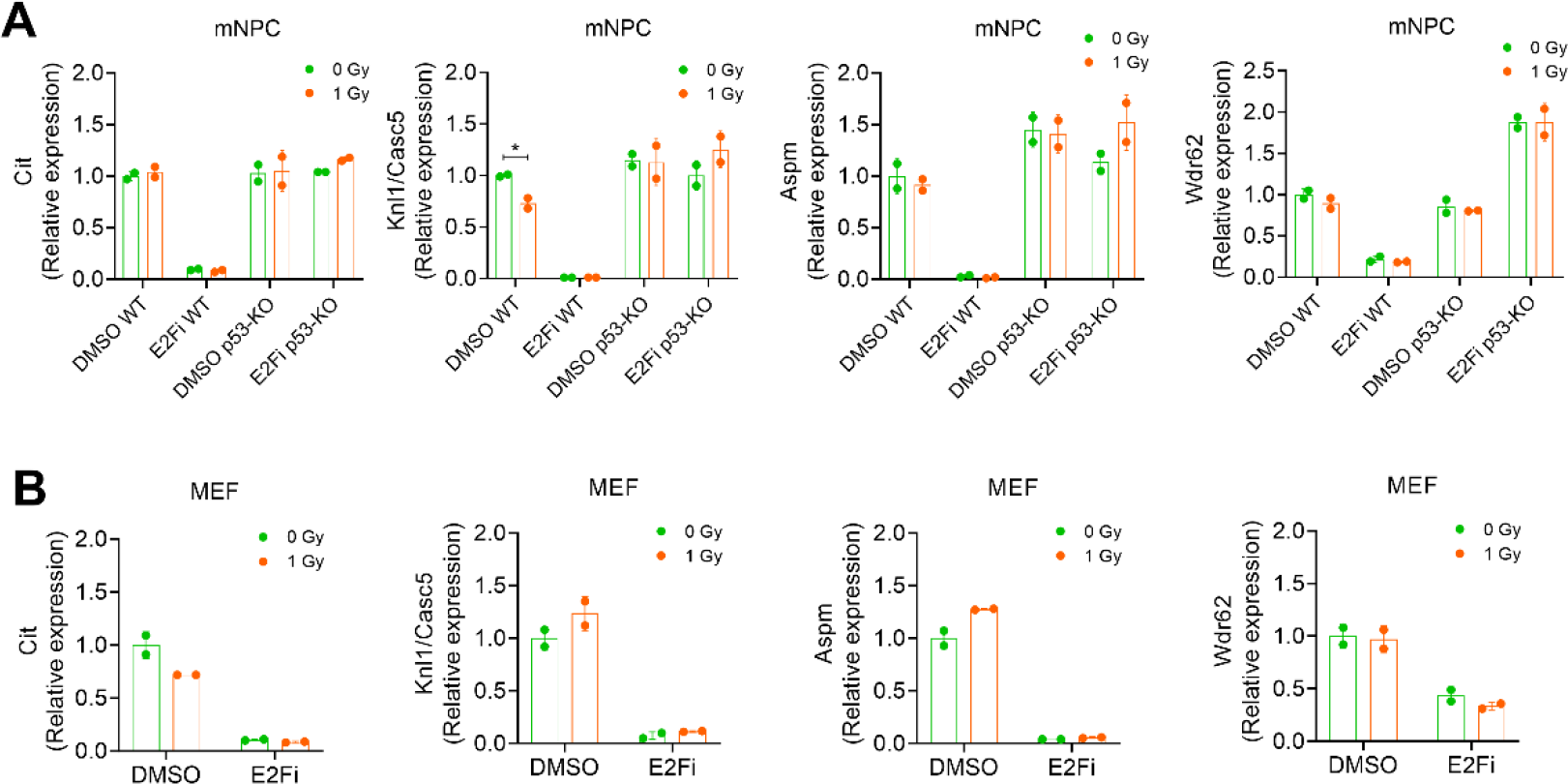
MCPH gene expression depends on p53-E2F in mouse NPCs and MEFs, but is not regulated by irradiation. (A) MCPH gene expression is reduced after treatment of wild-type primary mouse NPCs with an E2F inhibitor, but not after irradiation. E2F inhibition in p53 knockout mouse NPCs does not reduce MCPH gene expression. (B) MCPH gene expression is reduced after treatment of mouse embryonic fibroblasts with an E2F inhibitor, but not after irradiation. Student’s t-test was used, *p< 0.03.

## Notes

https://www.ebi.ac.uk/biostudies/arrayexpress

### Competing Interest Statement

The authors have declared no competing interest.

## References

1. Xing L, Wilsch-Bräuninger M, Huttner WB. How neural stem cells contribute to neocortex development. Biochem Soc Trans. 2021;49(5):1997–2006. doi:10.1042/BST20200923

2. O’Driscoll M, Jeggo PA. The role of the DNA damage response pathways in brain development and microcephaly: Insight from human disorders. DNA Repair (Amst). 2008;7(7):1039–1050. doi:10.1016/j.dnarep.2008.03.018

3. Pilaz LJ, McMahon JJ, Miller EE, Lennox AL, Suzuki A, Salmon E, Silver DL. Prolonged Mitosis of Neural Progenitors Alters Cell Fate in the Developing Brain. Neuron. 2016;89(1):83–99. doi:10.1016/j.neuron.2015.12.007

4. Phan TP, Holland AJ. Time is of the essence: the molecular mechanisms of primary microcephaly. Genes Dev. 2021;35(23-24):1551–1578. doi:10.1101/gad.348866.121

5. Jayaraman D, Bae BI, Walsh CA. The Genetics of Primary Microcephaly. Annu Rev Genomics Hum Genet. 2018;19(1):177–200. doi:10.1146/annurev-genom-083117-021441

6. Online Mendelian Inheritance in Man, OMIM®. Johns Hopkins University, Baltimore, MD. MIM Number: 251200. URL: https://www.omim.org/entry/251200.

7. Farcy S, Hachour H, Bahi-Buisson N, Passemard S. Genetic Primary Microcephalies: When Centrosome Dysfunction Dictates Brain and Body Size. Cells. 2023;12(13):1807. doi:10.3390/cells12131807

8. Ribeiro JH, Altinisik N, Rajan N, Verslegers M, Baatout S, Gopalakrishnan J, Quintens R. DNA damage and repair: underlying mechanisms leading to microcephaly. Front Cell Dev Biol. 2023;11(October):1–20. doi:10.3389/fcell.2023.1268565

9. Iegiani G, Ferraro A, Pallavicini G, Di Cunto F. The impact of TP53 activation and apoptosis in primary hereditary microcephaly. Front Neurosci. 2023;17. doi:10.3389/fnins.2023.1220010

10. McKinnon PJ. DNA repair deficiency and neurological disease. Nat Rev Neurosci. 2009;10(2):100–112. doi:10.1038/nrn2559

11. Otake M, Schull WJ. Review: Radiation-related brain damage and growth retardation among the prenatally exposed atomic bomb survivors. Int J Radiat Biol. 1998;74(2):159–171. doi:10.1080/095530098141555

12. Hammack C, Ogden SC, Madden JC, Medina A, Xu C, Phillips E, Son Y, Cone A, Giovinazzi S, Didier RA, Gilbert DM, Song H, Ming G, Wen Z, Brinton MA, Gunjan A, Tang H. Zika Virus Infection Induces DNA Damage Response in Human Neural Progenitors That Enhances Viral Replication. Heise MT, ed. J Virol. 2019;93(20):1–20. doi:10.1128/JVI.00638-19

13. Dunn K, Yoshimaru H, Otake M, Annegers JF, Schull WJ. Prenatal exposure to ionizing radiation and subsequent development of seizures. Am J Epidemiol. 1990;131(1):114–123. doi:10.1093/oxfordjournals.aje.a115464

14. McKinnon PJ. Maintaining genome stability in the nervous system. Nat Neurosci. 2013;16(11):1523–1529. doi:10.1038/nn.3537

15. Lakin ND, Jackson SP. Regulation of p53 in response to DNA damage. Oncogene. 1999;18(53):7644–7655. doi:10.1038/sj.onc.1203015

16. Vousden KH, Prives C. Blinded by the Light: The Growing Complexity of p53. Cell. 2009;137(3):413–431. doi:10.1016/j.cell.2009.04.037

17. Martins S, Erichsen L, Datsi A, Wruck W, Goering W, Chatzantonaki E, de Amorim VCM, Rossi A, Chrzanowska KH, Adjaye J. Impaired p53-Mediated DNA Damage Response Contributes to Microcephaly in Nijmegen Breakage Syndrome Patient-Derived Cerebral Organoids. Cells. 2022;11(5):802. doi:10.3390/cells11050802

18. Engeland K. Cell cycle arrest through indirect transcriptional repression by p53: I have a DREAM. Cell Death Differ. 2018;25(1):114–132. doi:10.1038/cdd.2017.172

19. Lang L, Pettkó-Szandtner A, Tunçay Elbaşı H, Takatsuka H, Nomoto Y, Zaki A, Dorokhov S, De Jaeger G, Eeckhout D, Ito M, Magyar Z, Bögre L, Heese M, Schnittger A. The DREAM complex represses growth in response to DNA damage in Arabidopsis. Life Sci Alliance. 2021;4(12):e202101141. doi:10.26508/lsa.202101141

20. Sadasivam S, DeCaprio JA. The DREAM complex: master coordinator of cell cycle-dependent gene expression. Nat Rev Cancer. 2013;13(8):585–595. doi:10.1038/nrc3556

21. Shimada M, Matsuzaki F, Kato A, Kobayashi J, Matsumoto T, Komatsu K. Induction of Excess Centrosomes in Neural Progenitor Cells during the Development of Radiation-Induced Microcephaly. Klymkowsky M, ed. PLoS One. 2016;11(7):e0158236. doi:10.1371/journal.pone.0158236

22. Verreet T, Rangarajan JR, Quintens R, Verslegers M, Lo AC, Govaerts K, Neefs M, Leysen L, Baatout S, Maes F, Himmelreich U, D’Hooge R, Moons L, Benotmane MA. Persistent Impact of In utero Irradiation on Mouse Brain Structure and Function Characterized by MR Imaging and Behavioral Analysis. Front Behav Neurosci. 2016;10(83):1–18. doi:10.3389/fnbeh.2016.00083

23. Mfossa ACM, Verslegers M, Verreet T, Fida H bin, Mysara M, Van IJcken W, De Vos W, Moons L, Baatout S, Benotmane M, Huylebroeck D, Quintens R. p53 drives premature neuronal differentiation in response to radiation-induced DNA damage during early neurogenesis. *bioRxiv*. Published online 2020:2020.06.26.171132. doi:10.1101/2020.06.26.171132

24. Saha S, Woodbine L, Haines J, Coster M, Ricket N, Barazzuol L, Ainsbury E, Sienkiewicz Z, Jeggo P. Increased apoptosis and DNA double-strand breaks in the embryonic mouse brain in response to very low-dose X-rays but not 50 Hz magnetic fields. J R Soc Interface. 2014;11(100):20140783. doi:10.1098/rsif.2014.0783

25. Roque T, Haton C, Etienne O, Chicheportiche A, Rousseau L, Martin L, Mouthon M, Boussin FD. Lack of a p21 waf1/cip -Dependent G1/S Checkpoint in Neural Stem and Progenitor Cells After DNA Damage In Vivo. Stem Cells. 2012;30(3):537–547. doi:10.1002/stem.1010

26. Lee Y, Chong MJ, McKinnon PJ. Ataxia Telangiectasia Mutated -Dependent Apoptosis after Genotoxic Stress in the Developing Nervous System Is Determined by Cellular Differentiation Status. J Neurosci. 2001;21(17):6687–6693. doi:10.1523/JNEUROSCI.21-17-06687.2001

27. Gatz SA, Ju L, Gruber R, Hoffmann E, Carr AM, Wang ZQ, Liu C, Jeggo PA. Requirement for DNA Ligase IV during Embryonic Neuronal Development. J Neurosci. 2011;31(27):10088–10100. doi:10.1523/JNEUROSCI.1324-11.2011

28. Williams SE, Garcia I, Crowther AJ, Li S, Stewart A, Liu H, Lough KJ, O’Neill S, Veleta K, Oyarzabal EA, Merrill JR, Shih YYI, Gershon TR. Aspm sustains postnatal cerebellar neurogenesis and medulloblastoma growth. Development. 2015;142(22):3921–3932. doi:10.1242/dev.124271

29. Gabriel E, Wason A, Ramani A, Gooi LM, Keller P, Pozniakovsky A, Poser I, Noack F, Telugu NS, Calegari F, Šarić T, Hescheler J, Hyman AA, Gottardo M, Callaini G, Alkuraya FS, Gopalakrishnan J. CPAP promotes timely cilium disassembly to maintain neural progenitor pool. EMBO J. 2016;35(8):803–819. doi:10.15252/embj.201593679

30. Verheyde J, de Saint-Georges L, Leyns L, Benotmane MA. The Role of Trp53 in the Transcriptional Response to Ionizing Radiation in the Developing Brain. DNA Res. 2006;13(2):65–75. doi:10.1093/dnares/dsi028

31. Quintens R, Verreet T, Janssen A, Neefs M, Leysen L, Michaux A, Verslegers M, Samari N, Pani G, Verheyde J, Baatout S, Benotmane MA. Identification of novel radiation- induced p53-dependent transcripts extensively regulated during mouse brain development. Biol Open. 2015;4(3):331–344. doi:10.1242/bio.20149969

32. Verreet T, Quintens R, Van Dam D, Verslegers M, Tanori M, Casciati A, Neefs M, Leysen L, Michaux A, Janssen A, D’Agostino E, Vande Velde G, Baatout S, Moons L, Pazzaglia S, Saran A, Himmelreich U, De Deyn PP, Benotmane MA. A multidisciplinary approach unravels early and persistent effects of X-ray exposure at the onset of prenatal neurogenesis. J Neurodev Disord. 2015;7(1):3. doi:10.1186/1866-1955-7-3

33. Bowen ME, Attardi LD. The role of p53 in developmental syndromes. J Mol Cell Biol. 2019;11(3):200–211. doi:10.1093/jmcb/mjy087

34. Gabriel E, Ramani A, Altinisik N, Gopalakrishnan J. Human Brain Organoids to Decode Mechanisms of Microcephaly. Front Cell Neurosci. 2020;14(May):1–13. doi:10.3389/fncel.2020.00115

35. Pulvers JN, Bryk J, Fish JL, Wilsch-Bräuninger M, Arai Y, Schreier D, Naumann R, Helppi J, Habermann B, Vogt J, Nitsch R, Tóth A, Enard W, Pääbo S, Huttner WB. Mutations in mouse Aspm (abnormal spindle-like microcephaly associated) cause not only microcephaly but also major defects in the germline. Proc Natl Acad Sci. 2010;107(38):16595–16600. doi:10.1073/pnas.1010494107

36. Jayaraman D, Kodani A, Gonzalez DM, Mancias JD, Mochida GH, Vagnoni C, Johnson J, Krogan N, Harper JW, Reiter JF, Yu TW, Bae B il, Walsh CA. Microcephaly Proteins Wdr62 and Aspm Define a Mother Centriole Complex Regulating Centriole Biogenesis, Apical Complex, and Cell Fate. Neuron. 2016;92(4):813–828. doi:10.1016/j.neuron.2016.09.056

37. Barrera JA, Kao LR, Hammer RE, Seemann J, Fuchs JL, Megraw TL. CDK5RAP2 Regulates Centriole Engagement and Cohesion in Mice. Dev Cell. 2010;18(6):913–926. doi:10.1016/j.devcel.2010.05.017

38. Lizarraga SB, Margossian SP, Harris MH, Campagna DR, Han AP, Blevins S, Mudbhary R, Barker JE, Walsh CA, Fleming MD. Cdk5rap2 regulates centrosome function and chromosome segregation in neuronal progenitors. Development. 2010;137(11):1907-1917. doi:10.1242/dev.040410

39. Verreet T, Verslegers M, Quintens R, Baatout S, Benotmane MA. Current Evidence for Developmental, Structural, and Functional Brain Defects following Prenatal Radiation Exposure. Neural Plast. 2016;2016(1243527):1–17. doi:10.1155/2016/1243527

40. Oyefeso FA, Goldberg G, Opoku NYPS, Vazquez M, Bertucci A, Chen Z, Wang C, Muotri AR, Pecaut MJ. Effects of acute low-moderate dose ionizing radiation to human brain organoids. Kanungo J, ed. PLoS One. 2023;18(5):e0282958. doi:10.1371/journal.pone.0282958

41. Bojcevski J, Wang C, Liu H, Abdollahi A, Dokic I. Assessment of Normal Tissue Radiosensitivity by Evaluating DNA Damage and Repair Kinetics in Human Brain Organoids. Int J Mol Sci. 2021;22(24):13195. doi:10.3390/ijms222413195

42. Van Assche IA, Van Calsteren K, Lemiere J, Hohmann J, Blommaert J, Huis in ’t Veld EA, Cardonick E, LeJeune C, Ottevanger NPB, Witteveen EPO, van Grotel M, van den Heuvel-Eibrink MM, Lagae L, Lambrecht M, Amant F. Long-term neurocognitive, psychosocial, and physical outcomes after prenatal exposure to radiotherapy: a multicentre cohort study of the International Network on Cancer, Infertility, and Pregnancy. Lancet Child Adolesc Heal. Published online April 2024. doi:10.1016/S2352-4642(24)00075-0

43. de Haan J, Verheecke M, Van Calsteren K, Van Calster B, Shmakov RG, Mhallem Gziri M, Halaska MJ, Fruscio R, Lok CAR, Boere IA, Zola P, Ottevanger PB, de Groot CJM, Peccatori FA, Dahl Steffensen K, Cardonick EH, Polushkina E, Rob L, Ceppi L, et al. Oncological management and obstetric and neonatal outcomes for women diagnosed with cancer during pregnancy: a 20-year international cohort study of 1170 patients. Lancet Oncol. 2018;19(3):337–346. doi:10.1016/S1470-2045(18)30059-7

44. Blommaert J, De Saint-Hubert M, Depuydt T, Oldehinkel E, Poortmans P, Amant F, Lambrecht M. Challenges and opportunities for proton therapy during pregnancy. Acta Obstet Gynecol Scand. 2024;103(4):767–774. doi:10.1111/aogs.14645

45. Sloan SA, Andersen J, Pașca AM, Birey F, Pașca SP. Generation and assembly of human brain region–specific three-dimensional cultures. Nat Protoc. 2018;13(9):2062–2085. doi:10.1038/s41596-018-0032-7

46. Iwata R, Vanderhaeghen P. Regulatory roles of mitochondria and metabolism in neurogenesis. Curr Opin Neurobiol. 2021;69:231–240. doi:10.1016/j.conb.2021.05.003

47. Jones FS, Meech R. Knockout of REST/NRSF shows that the protein is a potent repressor of neuronally expressed genes in non-neural tissues. BioEssays. 1999;21(5):372–376. doi:10.1002/(SICI)1521-1878(199905)21:5<372::AID-BIES3>3.0.CO;2-3

48. Dietrich N, Lerdrup M, Landt E, Agrawal-Singh S, Bak M, Tommerup N, Rappsilber J, Södersten E, Hansen K. REST–Mediated Recruitment of Polycomb Repressor Complexes in Mammalian Cells. Madhani HD, ed. PLoS Genet. 2012;8(3):e1002494. doi:10.1371/journal.pgen.1002494

49. Chambers SM, Fasano CA, Papapetrou EP, Tomishima M, Sadelain M, Studer L. Highly efficient neural conversion of human ES and iPS cells by dual inhibition of SMAD signaling. Nat Biotechnol. 2009;27(3):275–280. doi:10.1038/nbt.1529

50. Sah VP, Attardi LD, Mulligan GJ, Williams BO, Bronson RT, Jacks T. A subset of p53- deficient embryos exhibit exencephaly. Nat Genet. 1995;10(2):175–180. doi:10.1038/ng0695-175

51. Kim B, Koh Y, Do H, Ju Y, Choi J Bin, Cho G, Yoo HW, Lee BH, Han J, Park JE, Han YM. Aberrant Cortical Layer Development of Brain Organoids Derived from Noonan Syndrome-iPSCs. Int J Mol Sci. 2022;23(22):13861. doi:10.3390/ijms232213861

52. Cardoso-Moreira M, Halbert J, Valloton D, Velten B, Chen C, Shao Y, Liechti A, Ascenção K, Rummel C, Ovchinnikova S, Mazin P V., Xenarios I, Harshman K, Mort M, Cooper DN, Sandi C, Soares MJ, Ferreira PG, Afonso S, et al. Gene expression across mammalian organ development. Nature. 2019;571(7766):505-509. doi:10.1038/s41586-019-1338-5

53. Gordon A, Yoon SJ, Tran SS, Makinson CD, Park JY, Andersen J, Valencia AM, Horvath S, Xiao X, Huguenard JR, Pașca SP, Geschwind DH. Long-term maturation of human cortical organoids matches key early postnatal transitions. Nat Neurosci. 2021;24(3):331–342. doi:10.1038/s41593-021-00802-y

54. Das D, Li J, Cheng L, Franco S, Mahairaki V. Human Forebrain Organoids from Induced Pluripotent Stem Cells: A Novel Approach to Model Repair of Ionizing Radiation-Induced DNA Damage in Human Neurons. Radiat Res. 2020;194(2):191. doi:10.1667/RR15567.1

55. Jackson SP, Bartek J. The DNA-damage response in human biology and disease. Nature. 2009;461(7267):1071-1078. doi:10.1038/nature08467

56. Wang YH, Ho TLF, Hariharan A, Goh HC, Wong YL, Verkaik NS, Lee MY, Tam WL, van Gent DC, Venkitaraman AR, Sheetz MP, Lane DP. Rapid recruitment of p53 to DNA damage sites directs DNA repair choice and integrity. Proc Natl Acad Sci U S A. 2022;119(10):1–12. doi:10.1073/pnas.2113233119

57. Bates S, Vousden KH. Mechanisms of p53-mediated apoptosis. Cell Mol Life Sci C. 1999;55(1):28–37. doi:10.1007/s000180050267

58. Namba T, Nardelli J, Gressens P, Huttner WB. Metabolic Regulation of Neocortical Expansion in Development and Evolution. Neuron. 2021;109(3):408–419. doi:10.1016/j.neuron.2020.11.014

59. Spassky N, Meunier A. The development and functions of multiciliated epithelia. Nat Rev Mol Cell Biol. 2017;18(7):423–436. doi:10.1038/nrm.2017.21

60. Olstad EW, Ringers C, Hansen JN, Wens A, Brandt C, Wachten D, Yaksi E, Jurisch-Yaksi N. Ciliary Beating Compartmentalizes Cerebrospinal Fluid Flow in the Brain and Regulates Ventricular Development. Curr Biol. 2019;29(2):229–241.e6. doi:10.1016/j.cub.2018.11.059

61. Parichha A, Suresh V, Chatterjee M, Kshirsagar A, Ben-Reuven L, Olender T, Taketo MM, Radosevic V, Bobic-Rasonja M, Trnski S, Holtzman MJ, Jovanov-Milosevic N, Reiner O, Tole S. Constitutive activation of canonical Wnt signaling disrupts choroid plexus epithelial fate. Nat Commun. 2022;13(1):633. doi:10.1038/s41467-021-27602-z

62. Klingler E, Francis F, Jabaudon D, Cappello S. Mapping the molecular and cellular complexity of cortical malformations. Science (80-). 2021;371(6527). doi:10.1126/science.aba4517

63. Zhang J, Graham S, Tello A, Liu H, White MF. Multiple nucleic acid cleavage modes in divergent type III CRISPR systems. Nucleic Acids Res. 2016;44(4):1789–1799. doi:10.1093/nar/gkw020

64. Tang H, Hammack C, Ogden SC, Wen Z, Qian X. Zika Virus Infects Human Cortical Neural Precursors Attenuates Growth 2016 Tang. Cell Stem Cell. 2017;18(5):587–590. doi:10.1016/j.stem.2016.02.016.Zika

65. Kim J, Lee S, Lee J, Park JC, Kim KH, Ko JM, Park SH, Kim SK, Mook-Jung I, Lee JY. Neurotoxicity of phenylalanine on human iPSC-derived cerebral organoids. Mol Genet Metab. 2022;136(2):132–144. doi:10.1016/j.ymgme.2022.04.005

66. Moreau MX, Saillour Y, Elorriaga V, Bouloudi B, Delberghe E, Deutsch Guerrero T, Ochandorena-Saa A, Maeso-Alonso L, Marques MM, Marin MC, Spassky N, Pierani A, Causeret F. Repurposing of the multiciliation gene regulatory network in fate specification of Cajal-Retzius neurons. Dev Cell. 2023;58(15):1365–1382.e6. doi:10.1016/j.devcel.2023.05.011

67. Zaqout S, Morris-Rosendahl D, Kaindl A. Autosomal Recessive Primary Microcephaly (MCPH): An Update. Neuropediatrics. 2017;48(03):135–142. doi:10.1055/s-0037-1601448

68. Fujimori A, Yaoi T, Ogi H, Wang B, Suetomi K, Sekine E, Yu D, Kato T, Takahashi S, Okayasu R, Itoh K, Fushiki S. Ionizing radiation downregulates ASPM, a gene responsible for microcephaly in humans. Biochem Biophys Res Commun. 2008;369(3):953–957. doi:10.1016/j.bbrc.2008.02.149

69. Bianchi FT, Tocco C, Pallavicini G, Liu Y, Vernì F, Merigliano C, Bonaccorsi S, El- Assawy N, Priano L, Gai M, Berto GE, Chiotto AMA, Sgrò F, Caramello A, Tasca L, Ala U, Neri F, Oliviero S, Mauro A, et al. Citron Kinase Deficiency Leads to Chromosomal Instability and TP53-Sensitive Microcephaly. Cell Rep. 2017;18(7):1674–1686. doi:10.1016/j.celrep.2017.01.054

70. Shi L, Qalieh A, Lam MM, Keil JM, Kwan KY. Robust elimination of genome-damaged cells safeguards against brain somatic aneuploidy following Knl1 deletion. Nat Commun. 2019;10(1):2588. doi:10.1038/s41467-019-10411-w

71. Quintens R. Convergence and divergence between the transcriptional responses to Zika virus infection and prenatal irradiation. Cell Death Dis. 2017;8(3):e2672–e2672. doi:10.1038/cddis.2017.109

72. El Ghouzzi V, Bianchi FT, Molineris I, Mounce BC, Berto GE, Rak M, Lebon S, Aubry L, Tocco C, Gai M, Chiotto AMA, Sgrò F, Pallavicini G, Simon-Loriere E, Passemard S, Vignuzzi M, Gressens P, Di Cunto F. ZIKA virus elicits P53 activation and genotoxic stress in human neural progenitors similar to mutations involved in severe forms of genetic microcephaly and p53. Cell Death Dis. 2016;7(10):e2440–e2440. doi:10.1038/cddis.2016.266

73. Uxa S, Bernhart SH, Mages CFS, Fischer M, Kohler R, Hoffmann S, Stadler PF, Engeland K, Müller GA. DREAM and RB cooperate to induce gene repression and cell-cycle arrest in response to p53 activation. Nucleic Acids Res. 2019;47(17):9087–9103. doi:10.1093/nar/gkz635

74. Zou Z, Ohta T, Miura F, Oki S. ChIP-Atlas 2021 update: a data-mining suite for exploring epigenomic landscapes by fully integrating ChIP-seq, ATAC-seq and Bisulfite-seq data. Nucleic Acids Res. 2022;50(W1):W175-W182. doi:10.1093/nar/gkac199

75. Fischer M. Conservation and divergence of the p53 gene regulatory network between mice and humans. Oncogene. 2019;38(21):4095–4109. doi:10.1038/s41388-019-0706-9

76. Fischer M, Schwarz R, Riege K, DeCaprio JA, Hoffmann S. TargetGeneReg 2.0: a comprehensive web-atlas for p53, p63, and cell cycle-dependent gene regulation. NAR Cancer. 2022;4(1):zcac009. doi:10.1093/narcan/zcac009

77. Rakotopare J, Lejour V, Duval C, Eldawra E, Escoffier H, Toledo F. A systematic approach identifies p53-DREAM pathway target genes associated with blood or brain abnormalities. Dis Model Mech. 2023;16(10). doi:10.1242/dmm.050376

78. Lancaster MA, Renner M, Martin CA, Wenzel D, Bicknell LS, Hurles ME, Homfray T, Penninger JM, Jackson AP, Knoblich JA. Cerebral organoids model human brain development and microcephaly. Nature. 2013;501(7467):373-379. doi:10.1038/nature12517

79. Kang Y, Zhou Y, Li Y, Han Y, Xu J, Niu W, Li Z, Liu S, Feng H, Huang W, Duan R, Xu T, Raj N, Zhang F, Dou J, Xu C, Wu H, Bassell GJ, Warren ST, et al. A human forebrain organoid model of fragile X syndrome exhibits altered neurogenesis and highlights new treatment strategies. Nat Neurosci. 2021;24(10):1377–1391. doi:10.1038/s41593-021-00913-6

80. Chen M, Lee HK, Moo L, Hanlon E, Stein T, Xia W. Common proteomic profiles of induced pluripotent stem cell-derived three-dimensional neurons and brain tissue from Alzheimer patients. J Proteomics. 2018;182:21–33. doi:10.1016/j.jprot.2018.04.032

81. Benito-Kwiecinski S, Lancaster MA. Brain Organoids: Human Neurodevelopment in a Dish. Cold Spring Harb Perspect Biol. 2020;12(8):a035709. doi:10.1101/cshperspect.a035709

82. Camp JG, Badsha F, Florio M, Kanton S, Gerber T, Wilsch-Bräuninger M, Lewitus E, Sykes A, Hevers W, Lancaster M, Knoblich JA, Lachmann R, Pääbo S, Huttner WB, Treutlein B. Human cerebral organoids recapitulate gene expression programs of fetal neocortex development. Proc Natl Acad Sci. 2015;112(51):15672–15677. doi:10.1073/pnas.1520760112

83. Fernandes S, Klein D, Marchetto MC. Unraveling Human Brain Development and Evolution Using Organoid Models. Front Cell Dev Biol. 2021;9. doi:10.3389/fcell.2021.737429

84. Jaylet T, Quintens R, Benotmane MA, Luukkonen J, Tanaka IB, Ibanez C, Durand C, Sachana M, Azimzadeh O, Adam-Guillermin C, Tollefsen KE, Laurent O, Audouze K, Armant O. Development of an adverse outcome pathway for radiation-induced microcephaly via expert consultation and machine learning. Int J Radiat Biol. 2022;98(12):1752–1762. doi:10.1080/09553002.2022.2110312

85. Jaylet T, Quintens R, Armant O, Audouze K. An integrative systems biology strategy to support the development of adverse outcome pathways (AOPs): a case study on radiation- induced microcephaly. Front Cell Dev Biol. 2023;11. doi:10.3389/fcell.2023.1197204

86. Otake M, Schull WJ. Radiation-related Small Head Sizes among Prenatally Exposed A- bomb Survivors. Int J Radiat Biol. 1993;63(2):255–270. doi:10.1080/09553009314550341

87. Rakotomamonjy J, Rylaarsdam L, Fares-Taie L, McDermott S, Davies D, Yang G, Fagbemi F, Epstein M, Fairbanks-Santana M, Rozet JM, Guemez-Gamboa A. PCDH12 loss results in premature neuronal differentiation and impeded migration in a cortical organoid model. Cell Rep. 2023;42(8):112845. doi:10.1016/j.celrep.2023.112845

88. Lee Y, Katyal S, Downing SM, Zhao J, Russell HR, McKinnon PJ. Neurogenesis requires TopBP1 to prevent catastrophic replicative DNA damage in early progenitors. Nat Neurosci. 2012;15(6):819–826. doi:10.1038/nn.3097

89. Kalogeropoulou A, Lygerou Z, Taraviras S. Cortical Development and Brain Malformations: Insights From the Differential Regulation of Early Events of DNA Replication. Front Cell Dev Biol. 2019;7(MAR):1–9. doi:10.3389/fcell.2019.00029

90. Durante M, Bender T, Schickel E, Mayer M, Debus J, Grosshans D, Schroeder I. Aberrant choroid plexus formation in human cerebral organoids exposed to radiation. Res Sq. 2023;Oct 17(rs.3.rs-3445801).

91. Fischer M, Steiner L, Engeland K. The transcription factor p53: Not a repressor, solely an activator. Cell Cycle. 2014;13(19):3037–3058. doi:10.4161/15384101.2014.949083

92. Fischer M. Census and evaluation of p53 target genes. Oncogene. 2017;36(28):3943–3956. doi:10.1038/onc.2016.502

93. Ma Y, Kurtyka CA, Boyapalle S, Sung SS, Lawrence H, Guida W, Cress WD. A Small- Molecule E2F Inhibitor Blocks Growth in a Melanoma Culture Model. Cancer Res. 2008;68(15):6292–6299. doi:10.1158/0008-5472.CAN-08-0121

94. Engeland K. Cell cycle regulation: p53-p21-RB signaling. Cell Death Differ. 2022;29(5):946–960. doi:10.1038/s41418-022-00988-z

95. Yoon J, Grinchuk O V., Tirado-Magallanes R, Ngian ZK, Tay EXY, Chuah YH, Lee BWL, Feng J, Crasta KC, Ong CT, Benoukraf T, Ong DST. E2F and STAT3 provide transcriptional synergy for histone variant H2AZ activation to sustain glioblastoma chromatin accessibility and tumorigenicity. Cell Death Differ. 2022;29(7):1379–1394. doi:10.1038/s41418-021-00926-5

96. Fischer M, Grossmann P, Padi M, DeCaprio JA. Integration of TP53, DREAM, MMB-FOXM1 and RB-E2F target gene analyses identifies cell cycle gene regulatory networks. Nucleic Acids Res. 2016;44(13):6070-6086. doi:10.1093/nar/gkw523

97. Rayon T, Stamataki D, Perez-Carrasco R, Garcia-Perez L, Barrington C, Melchionda M, Exelby K, Lazaro J, Tybulewicz VLJ, Fisher EMC, Briscoe J. Species-specific pace of development is associated with differences in protein stability. Science (80-). 2020;369(6510). doi:10.1126/science.aba7667

98. Pallavicini G, Berto GE, Di Cunto F. Precision Revisited: Targeting Microcephaly Kinases in Brain Tumors. Int J Mol Sci. 2019;20(9):2098. doi:10.3390/ijms20092098

99. Iegiani G, Di Cunto F, Pallavicini G. Inhibiting microcephaly genes as alternative to microtubule targeting agents to treat brain tumors. Cell Death Dis. 2021;12(11):956. doi:10.1038/s41419-021-04259-6

100. Bhaduri A, Andrews MG, Mancia Leon W, Jung D, Shin D, Allen D, Jung D, Schmunk G, Haeussler M, Salma J, Pollen AA, Nowakowski TJ, Kriegstein AR. Cell stress in cortical organoids impairs molecular subtype specification. Nature. 2020;578(7793):142-148. doi:10.1038/s41586-020-1962-0

101. Zhao HH, Haddad G. Brain organoid protocols and limitations. Front Cell Neurosci. 2024;18. doi:10.3389/fncel.2024.1351734

102. Martins-Costa C, Pham VA, Sidhaye J, Novatchkova M, Wiegers A, Peer A, Möseneder P, Corsini NS, Knoblich JA. Morphogenesis and development of human telencephalic organoids in the absence and presence of exogenous extracellular matrix. EMBO J. 2023;42(22). doi:10.15252/embj.2022113213

103. Ewels PA, Peltzer A, Fillinger S, Patel H, Alneberg J, Wilm A, Garcia MU, Di Tommaso P, Nahnsen S. The nf-core framework for community-curated bioinformatics pipelines. Nat Biotechnol. 2020;38(3):276–278. doi:10.1038/s41587-020-0439-x

104. Jew B, Alvarez M, Rahmani E, Miao Z, Ko A, Garske KM, Sul JH, Pietiläinen KH, Pajukanta P, Halperin E. Accurate estimation of cell composition in bulk expression through robust integration of single-cell information. Nat Commun. 2020;11(1):1971. doi:10.1038/s41467-020-15816-6

105. Piron A, Szymczak F, Papadopoulou T, Alvelos MI, Defrance M, Lenaerts T, Eizirik DL, Cnop M. RedRibbon: A new rank–rank hypergeometric overlap for gene and transcript expression signatures. Life Sci Alliance. 2024;7(2):e202302203. doi:10.26508/lsa.202302203

106. Kuleshov M V., Jones MR, Rouillard AD, Fernandez NF, Duan Q, Wang Z, Koplev S, Jenkins SL, Jagodnik KM, Lachmann A, McDermott MG, Monteiro CD, Gundersen GW, Ma’ayan A. Enrichr: a comprehensive gene set enrichment analysis web server 2016 update. Nucleic Acids Res. 2016;44(W1):W90–W97. doi:10.1093/nar/gkw377

107. Zhou Y, Zhou B, Pache L, Chang M, Khodabakhshi AH, Tanaseichuk O, Benner C, Chanda SK. Metascape provides a biologist-oriented resource for the analysis of systems- level datasets. Nat Commun. 2019;10(1):1523. doi:10.1038/s41467-019-09234-6

108. Shannon P, Markiel A, Ozier O, Baliga NS, Wang JT, Ramage D, Amin N, Schwikowski B, Ideker T. Cytoscape: A Software Environment for Integrated Models of Biomolecular Interaction Networks. Genome Res. 2003;13(11):2498–2504. doi:10.1101/gr.1239303

109. Robinson MD, McCarthy DJ, Smyth GK. edgeR : a Bioconductor package for differential expression analysis of digital gene expression data. Bioinformatics. 2010;26(1):139–140. doi:10.1093/bioinformatics/btp616

110. Robinson MD, Oshlack A. A scaling normalization method for differential expression analysis of RNA-seq data. Genome Biol. 2010;11(3):R25. doi:10.1186/gb-2010-11-3-r25

111. Lovering RC, Gaudet P, Acencio ML, Ignatchenko A, Jolma A, Fornes O, Kuiper M, Kulakovskiy I V., Lægreid A, Martin MJ, Logie C. A GO catalogue of human DNA- binding transcription factors. Biochim Biophys Acta - Gene Regul Mech. 2021;1864(11- 12):194765. doi:10.1016/j.bbagrm.2021.194765

112. Piñero J, Ramírez-Anguita JM, Saüch-Pitarch J, Ronzano F, Centeno E, Sanz F, Furlong LI. The DisGeNET knowledge platform for disease genomics: 2019 update. Nucleic Acids Res. Published online November 4, 2019. doi:10.1093/nar/gkz1021

113. Bonnet E, Calzone L, Michoel T. Integrative Multi-omics Module Network Inference with Lemon-Tree. Gardner PP, ed. PLOS Comput Biol. 2015;11(2):e1003983. doi:10.1371/journal.pcbi.1003983

114. Badia-i-Mompel P, Vélez Santiago J, Braunger J, Geiss C, Dimitrov D, Müller-Dott S, Taus P, Dugourd A, Holland CH, Ramirez Flores RO, Saez-Rodriguez J. decoupleR: ensemble of computational methods to infer biological activities from omics data. Kuijjer ML, ed. Bioinforma Adv. 2022;2(1). doi:10.1093/bioadv/vbac016

115. Müller-Dott S, Tsirvouli E, Vazquez M, Ramirez Flores RO, Badia-i-Mompel P, Fallegger R, Türei D, Lægreid A, Saez-Rodriguez J. Expanding the coverage of regulons from high- confidence prior knowledge for accurate estimation of transcription factor activities. Nucleic Acids Res. 2023;51(20):10934–10949. doi:10.1093/nar/gkad841

116. Aibar S, González-Blas CB, Moerman T, Huynh-Thu VA, Imrichova H, Hulselmans G, Rambow F, Marine JC, Geurts P, Aerts J, van den Oord J, Atak ZK, Wouters J, Aerts S. SCENIC: single-cell regulatory network inference and clustering. Nat Methods. 2017;14(11):1083–1086. doi:10.1038/nmeth.4463

117. Pfaffl MW. A new mathematical model for relative quantification in real-time RT-PCR. Nucleic Acids Res. 2001;29(9):45e–45. doi:10.1093/nar/29.9.e45

